# Dysbiosis personalizes fitness effect of antibiotic resistance in the mammalian gut

**DOI:** 10.1101/748897

**Authors:** Luís Leónidas Cardoso, Paulo Durão, Massimo Amicone, Isabel Gordo

**Affiliations:** Instituto Gulbenkian de Ciência, Oeiras, Portugal

## Abstract

The fitness cost of antibiotic resistance in the absence of antibiotics is crucial to the success of suspending antibiotics as a strategy to lower resistance. Here we show that after antibiotic treatment the cost of resistance within the complex ecosystem of the mammalian gut is personalized. Using mice as an *in vivo* model, we find that the fitness effect of the same resistant mutation can be deleterious in a host, but neutral or even beneficial in other hosts. Such antagonistic pleiotropy is shaped by the microbiota, as in germ-free mice resistance is consistently costly across all hosts. An eco-evolutionary model of competition for resources identifies a general mechanism underlying between host variation and predicts that the dynamics of compensatory evolution of resistant bacteria should be host specific, a prediction that was supported by experimental evolution *in vivo*. The microbiome of each human is close to unique and our results suggest that the short-term costs of resistance and its long-term within-host evolution will also be highly personalized, a finding that may contribute to the observed variable outcome of control therapies.

**One Sentence Summary:** Personalized Fitness of Resistance Mutations.

## INTRODUCTION

Antibiotic resistance (AR) is a growing challenge in the treatment of infectious diseases which are projected to become a burden worldwide in the coming decades^1^. The set of AR genes and AR mutations – called resistomes – is widespread in clinical^2,3^ and environmental^4,5^ settings, providing a reservoir that can further expand by horizontal gene transfer. Commensal bacteria can carry AR in healthy individuals and AR can persist in the human gut for years^6^.

Chromosomal encoded resistance mutations often map onto genes coding for essential cellular functions, such as transcription, translation, or cell-wall biogenesis (see e.g.^7,8^). Resistance tends to be highly epistatic and pleiotropic^9–11^ and typically entails fitness costs in the absence of antibiotics^12–15^. The existence of AR costs predicts that a susceptible strain should out-compete a resistant strain, and a decrease of resistance levels to a given antibiotic should occur if the use of that drug is halted in clinical settings. This strategy should be effective when the cost of resistance is high^16–19^, allowing for the elimination of the AR strain before evolutionary compensation for the cost of resistance occurs^8^. Thus, the efficacy of controlling the spread of AR by suspending the usage of an antibiotic is critically dependent on the relative fitness of resistant and sensitive genotypes in the absence of antibiotic.

The costs of AR are strongly influenced by the environment where bacteria grow, both in its abiotic (e.g. nutrient availability) and biotic (interactions with other cells) components^20–22^. Fitness costs of AR can also depend on the genetic background, including the presence of other resistances, at the level of the core and accessory genome^9,23,24^. Since the effects of AR mutations have often been measured under laboratory environments, which lack the multiple interactions likely to be important *in natura*, our understanding of how costly AR can actually be is currently limited. A few studies where pathogens^25–31^ were tested during *in vivo* colonization and infection suggest that fitness costs of AR are not always high in the context of bacterial colonization or virulence. Yet to the best of our knowledge, no study so far has analyzed the temporal dynamics of resistant strains colonizing the key ecosystem of the gut microbiota. In particular, it is currently unclear how the results from *in vitro* studies or in the context of invasive pathogens are informative about AR in gut commensal strains, which are by far the main colonizers of a natural complex ecosystem. Here, we performed *in vivo* competitive fitness assays, mathematical modeling and *in vivo* experimental evolution to unravel the fitness effects of AR in commensal *E. coli* colonizing its natural environment.

## RESULTS

### Competitive fitness of AR in the mouse gut

We focused on common resistance mutations to streptomycin - Str^R^ (*rpsL*^K43T^) and rifampicin- Rif^R^ (*rpoB*^H526Y^), and also studied double resistant clones - Str^R^Rif^R^ (*rpsL*^K43T^*rpoB*^H526Y^). These have been identified in many important pathogens, such as *Mycobacterium tuberculosis* and Salmonella, and also in pathogenic and commensal *E. coli*^32–34^.

To query how inter-species interactions, present in the natural ecosystem comprising the mammalian gut, influence the costs of AR, we performed competitive fitness assays in mice that have a complex microbiota (SPF mice). To mimic conditions where the rise of AR can occur, mice were given an antibiotic treatment – streptomycin - for a week (see **Fig. 1a** and **Methods**). Such treatment is known to cause perturbations in the microbiota species composition and also to break colonization resistant to *E. coli*^35^, thus increasing the probability that colonization by external strains occurs. To measure the *in vivo* fitness effects of AR and quantify their costs, which should occur in the absence of antibiotic, we removed the treatment for two days and then colonized the mice with susceptible and resistant *E. coli* strains (**Fig. 1a**). Previous studies suggest that streptomycin is quickly removed from mice^36^ and we have experimentally confirmed that streptomycin is absent 2 days after treatment is stopped, via a biological detection method in the fecal samples (the developed method has a threshold of detection of ≈2μg/ml) (**Supplementary Fig. 1**). In agreement, we see variation of costs even when the competition is between two strains that are resistant to the streptomycin (**Supplementary Fig. 2**).

**Figure 1.**
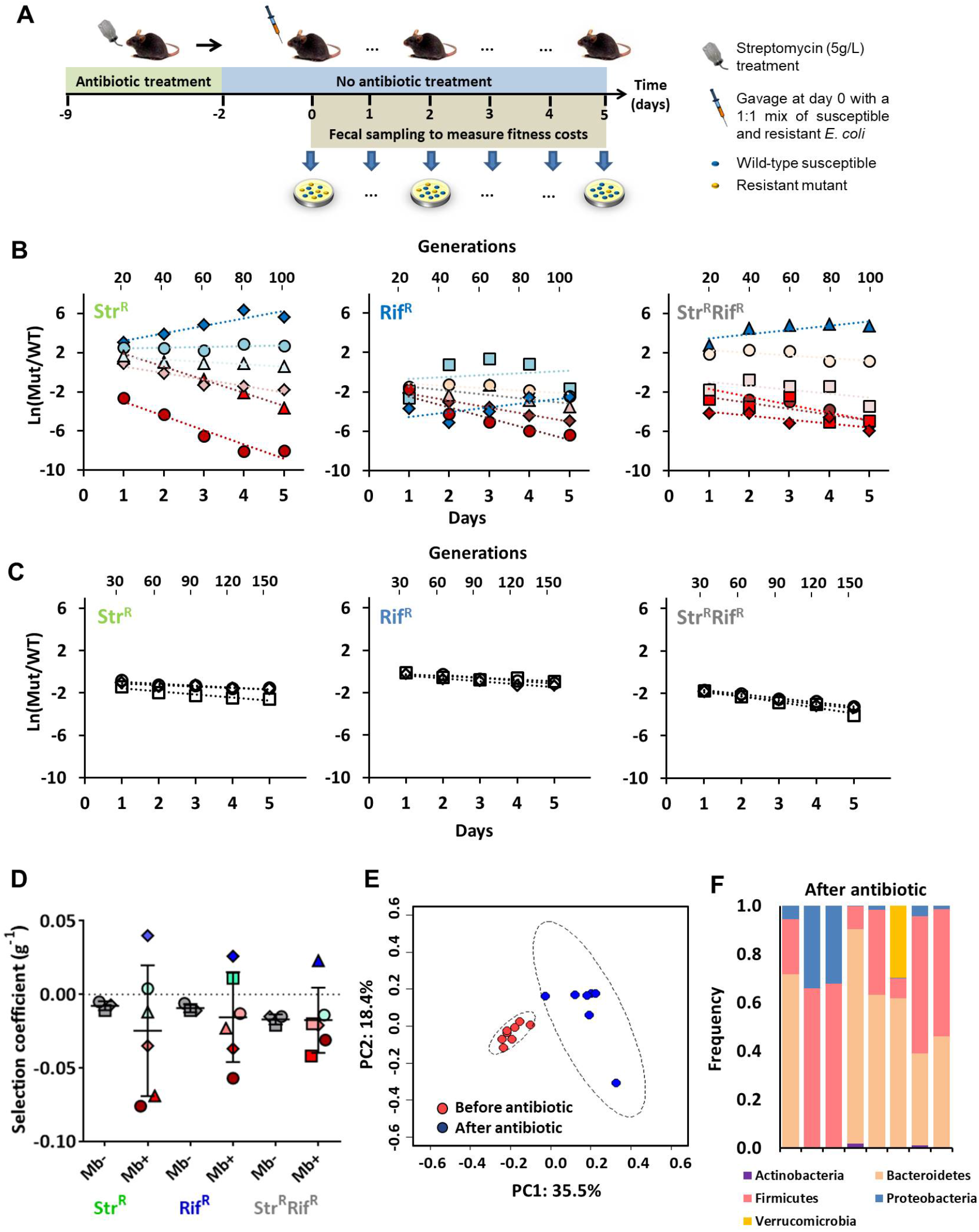
Effect of microbiota on the fitness costs of resistances. **(a)** Scheme of the experimental design to measure the fitness effect of AR *in vivo*. Mice with their natural microbiota were given a one-week course of streptomycin treatment, after which the antibiotic was removed from the water. Two days post-treatment mice were fed with a mixture of sensitive and AR *E. coli* strains, isogenic and marked with YFP and CFP respectively. The temporal dynamics of the AR frequency was estimated from plating of fecal samples daily. **(b,c)** The fitness effect of streptomycin resistance, coded by *rpsL*^*K43T*^ mutation (Str^R^), rifampicin resistance, coded by *rpoB*^*H526Y*^ mutation (Rif^R^), and the *rpsL*^*K43T*^*rpoB*^*H526Y*^ double mutant (Str^R^Rif^R^) under competition against a sensitive background in the presence of a diverse microbiota **(b)** and in the absence of inter-species interactions **(c)**. **(d)** Boxplot of the mean and variance of the fitness costs of resistance measured in mice mono-colonized and with a complex microbiota. **(e)** Microbiota beta diversity visualization by principal coordinate analysis (PCoA) based on Unweighted UniFrac distance before and after antibiotic treatment. Ellipses represent the standard deviation of point scores with a 95% confidence limit for each group (ANOSIM test, p < 0.05). **(f)** Microbiota composition as relative OTU abundance assayed by 16S rRNA amplicon sequencing and clustered at the phylum level (colored segments) in different mice after antibiotic treatment displaying the broader diversity across hosts observed in the PCoA.

For most of the competitions, the temporal dynamics of each of the resistant strains in each mouse, and hence, the fitness effects of AR within a host, were consistent with a constant selective effect throughout 5 days of colonization (**Fig. 1b**). However, a wide variation in the temporal dynamics of Log (AR/Susceptible) is observed between each mouse (**Fig. 1b**). Such variation is not the result of sampling noise but unveils host-specific fitness effects of AR. Remarkably AR caused a strong deleterious effect in a particular host, whereas in another host AR did not exhibit a significant cost (**Fig. 1b** and **Supplementary Table 1**). These results strongly suggest that the elimination of AR will likely take a very long time to occur, or may not occur at all, in certain hosts. Frequency dependent selection is unlikely to be the cause of the observed temporal variation in the frequency of AR between hosts, as the initial frequency of the resistant strain is not predictive of the resistance fate (**Fig. 1b)**. The occurrence of compensatory mutations, although possible, is also unlikely to explain the observed variation. Such events would have to be very common and also entail strong beneficial effects to influence the estimated fitness difference within the 5 day period studied. Compensatory mutations are also expected to take longer periods to be rise in frequency (see below) and their spread should lead to strong temporal variations in the frequency of AR strains within each mouse, causing significant deviations from a simple linear model, a pattern which was not observed. The data strongly suggests that constant selection against resistance occurs in a host, but selection for resistance can occur in another host during the 5-day co-colonization period. Such observation cannot be explained by the occurrence of back-mutations or by mutations that would render the bacteria sensitive to the antibiotic (**Supplementary Table 2**).

To investigate if the presence of a complex microbiota is an important contributor to the personalized fitness of AR, we performed co-colonization experiments in germ-free mice. Here the *in vivo* fitness costs of AR are solely derived from intra-strain competition in the gut and we find much lower variance between these hosts. Significant fitness costs of each resistant strain were estimated in this *in vivo* but simpler environment: 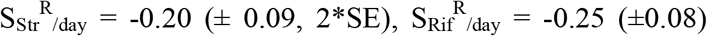 and 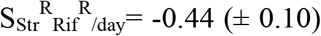 (**Fig. 1c** and **Supplementary Table 3-4**, corresponding to 1 to 2% cost per generation^37^, implying that AR should be eliminated within 50 to 100 generations, in the absence of antibiotics. The fitness effects of AR estimated *in vivo* are significantly different from those estimated *in vitro* (**Supplementary Fig. 3**). Indeed none of the commonly used laboratory environments provides a good predictor to the costs of single AR mutations, in the simplest *in vivo* system lacking a complex microbiota, nor of their combined effects (see **Supplemental Text** and **Supplemental Fig. 3**).

Having found that the fitness effects of AR are host-specific, we next asked about their effects at the population level. Taking the cohort of mice studied as a population, we find that AR is costly on average (**Fig. 1d**), although it is not significant in any of the cases 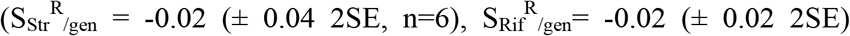 and 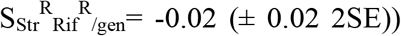. This indicates that all resistance strains would be difficult to eliminate at the host meta-population level.

Characterization of the gut microbiota composition of the cohorts of mice, through 16S rRNA sequencing, showed that antibiotic treatment both reduced the alpha diversity (p<0.001) and increased substantially the variation of the host microbiota (**Fig. 1e-f**). These results suggest that the personalized fitness effect of AR likely results from an interaction between the effect of AR and the microbial gut ecosystem.

### Modeling AR costs within a species rich ecosystem

To understand whether general properties of the microbiota could cause host-specific effects of mutations we turned to a theoretical model. If most prevalent interactions in the microbiota are competitive, as suggested by previous analysis^38^, we can use the MacArthur consumer-resource model, which only assumes competition. This framework is capable of explaining major diversity patterns of microbial communities^39^. The model was adapted to quantify the effect of a diverse microbiota on the relative fitness of an AR mutation (which is costly in the absence of other species) both analytically and numerically. This theoretical framework seems appropriate since the resistances studied affect core genes in bacterial metabolism and alter growth rates in different carbon sources (**Supplementary Fig. 4**). We assume that bacteria compete for a set of non-essential resources *S*_1_, …, *S*_*P*_ and each species is defined by their resource consumption rates 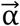 in a *P*-dimensional phenotypic space (**Fig. 2a-b**). We start by assuming that the species of the microbiota (*M*) initially satisfy the conditions for stable coexistence (**Supplementary Text, eq.3**). To quantify the fitness effect in this context, we assume that a mutant has a phenotypic difference from its parental wild type 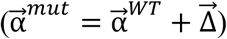 such that its overall fitness is impaired relative to the total amount of resources that the parent could consume 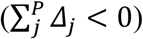. The selection coefficient in the presence of other species is then given by:

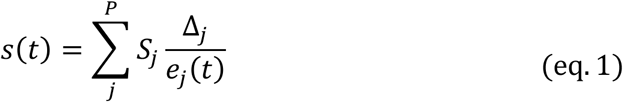

where 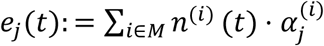 with *n*^(*i*)^ being the concentration of species *i*. The phenotypes modeled here can be thought of as enzymes dynamics, e_j_ (*t*), as they represent key functional units likely to be relevant in the competitive environment. Their abundance in the ecosystem (which is proportional to the density of the species) can vary over time, especially in the context of a strong perturbation, such as the antibiotic treatment in our experimental system. From the time-dependent form of selection in(eq. 1) one can deduce the following results: Firstly, at equilibrium (*e*_*j*_ = *S*_*j*_ ∀*j*), selection on the traits is additive, constant and independent of the microbiota composition (**Supplementary Text, eq.7**). However, the presence of a stable microbiota can amplify or buffer the cost of a mutation, according to its specific effect (**Fig. 2c, Supplementary Text, eq.8**). Furthermore, the probability that the cost is buffered increases with the ratio between the traits (**Fig. 2d**, **Supplementary Text, eq.8-9**). Secondly, when the microbiota ecosystem is pushed out of equilibrium via a perturbation (e.g. antibiotic treatment), the fate of a previously deleterious mutation can be significantly altered. Under such conditions, selection on the mutant becomes host-specific and can be negative, neutral or even positive in the short-term (**Supplementary Text, eq.10-11**).

**Figure 2.**
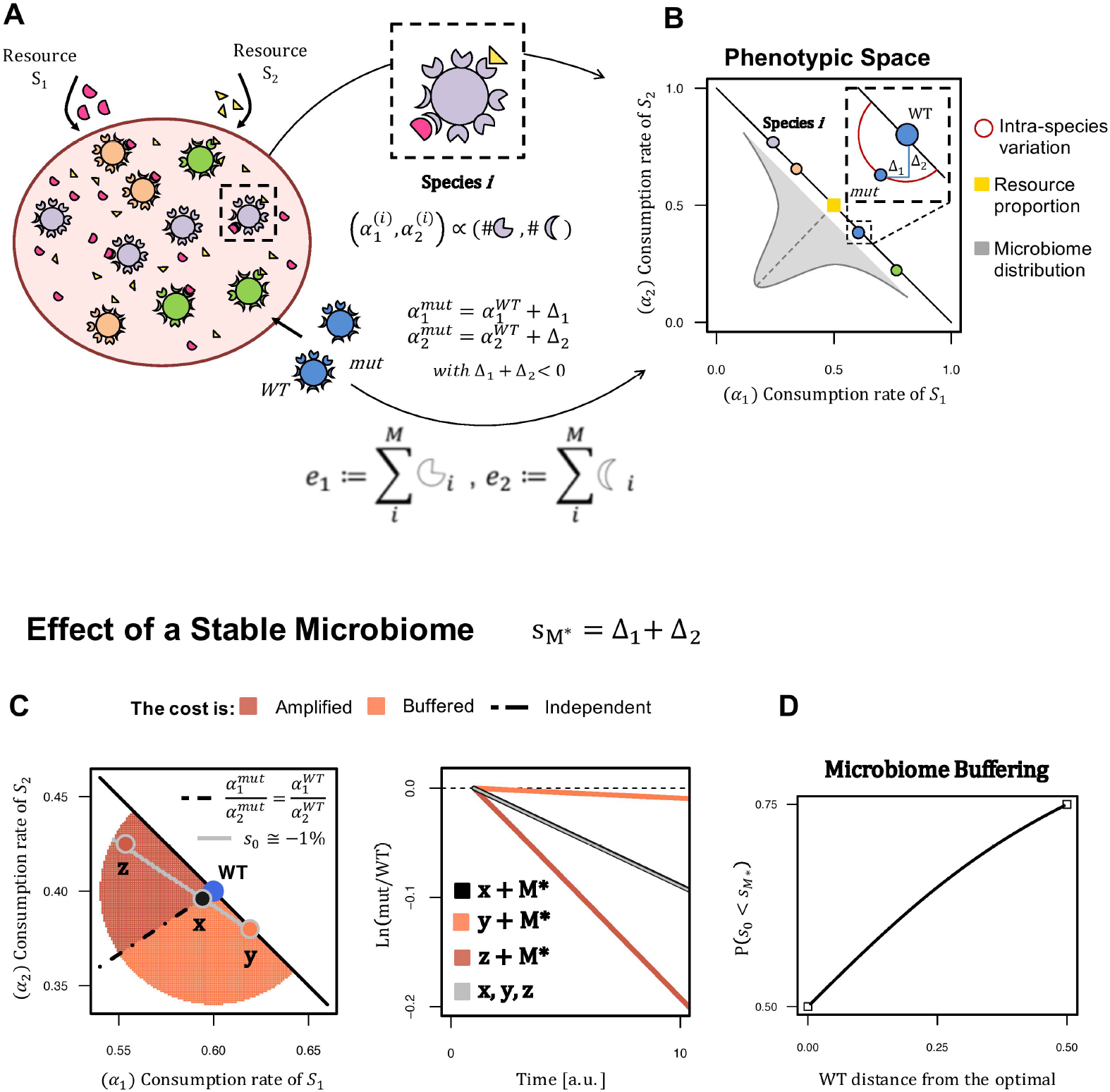
Multi-species ecological model of pleiotropic AR mutations and the effect of a stable microbiome. **(a)** Schematic of the model with two resources and multiple species. Each species *i* (represented by a given color) is characterized by its ability of consuming resources (*S*_1_ and *S*_2_), encoded by the traits 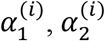 (represented by resource-specific shapes around each cell). **(b)** Species 2-D phenotypic space assuming a metabolic trade-off (species lie on the diagonal (see Supplementary Text)) to allow an equilibrium species rich state. **(c)** The relative fitness of a mutant in the presence of a stable microbiota (M), *s*_*M*\*_, is time-independent and independent of the specific composition of *M*. It however can be buffered or amplified by the microbiota according to the specific values of the mutation effect (Δ_1_, Δ_2_): when the trait ratio remains unchanged (e.g. Mutant x), *s* is not affected by other species, otherwise the cost can increase (e.g. Mutant z) or be buffered by the microbiota (e.g. Mutant y). **(d)** The probability of buffering increases with the distance of the WT to the theoretical optimal (yellow square in panel b, **Supplementary Text**, **eq8**).

We performed numerical simulations (see **Supplementary Text**) for the case of 2-resources to illustrate how the time dynamics predicted by the model may explain the experimental patterns in **Fig. 1**. The Ln(mutant/susceptible) varies in time and depends on the specific mutant (see **Fig. 3**). If a mutation changes two traits but not their ratio, its fitness cost is independent of the microbiota composition (**Fig. 3a-b**, Mutant *x*). For a mutation that causes an increase on one trait but a decrease on the other, the functional content of the ecosystem determines which trait is beneficial or detrimental and consequently, determines if the mutant is selected for or against (**Fig. 3a-b**, Mutants *y* and *z* have opposite fitness effects). Thus the model predicts variable fitness effects across hosts and reveals how a pleiotropy-dependent mechanism characteristic of AR mutants, can lead to their increase in frequency in the absence of antibiotics (**Supplementary Text, eq.11**). Importantly, at longer time scales, as the whole microbial ecosystem approaches equilibrium, the fitness effects converge towards a negative value (**Fig. 3c**), which will eventually become constant across all individuals (**Supplementary Text, eq.7)**. These results indicate that an AR mutation, which affects competition for resources, should exhibit a host-specific fitness effect during the initial days of competition (**Fig. 1b**), and predict that the AR cost should become host-independent once the microbiota reach equilibrium within a host. Importantly, since mode and time for equilibrium to occur are microbiota-dependent (**Supplementary Figure 5**), one can further predict that the selective pressure for compensatory mutations should be different across individuals. Thus, the dynamics of compensation for AR costs should be time-dependent with compensatory mutations appearing sooner in some hosts and later in others.

**Figure 3.**
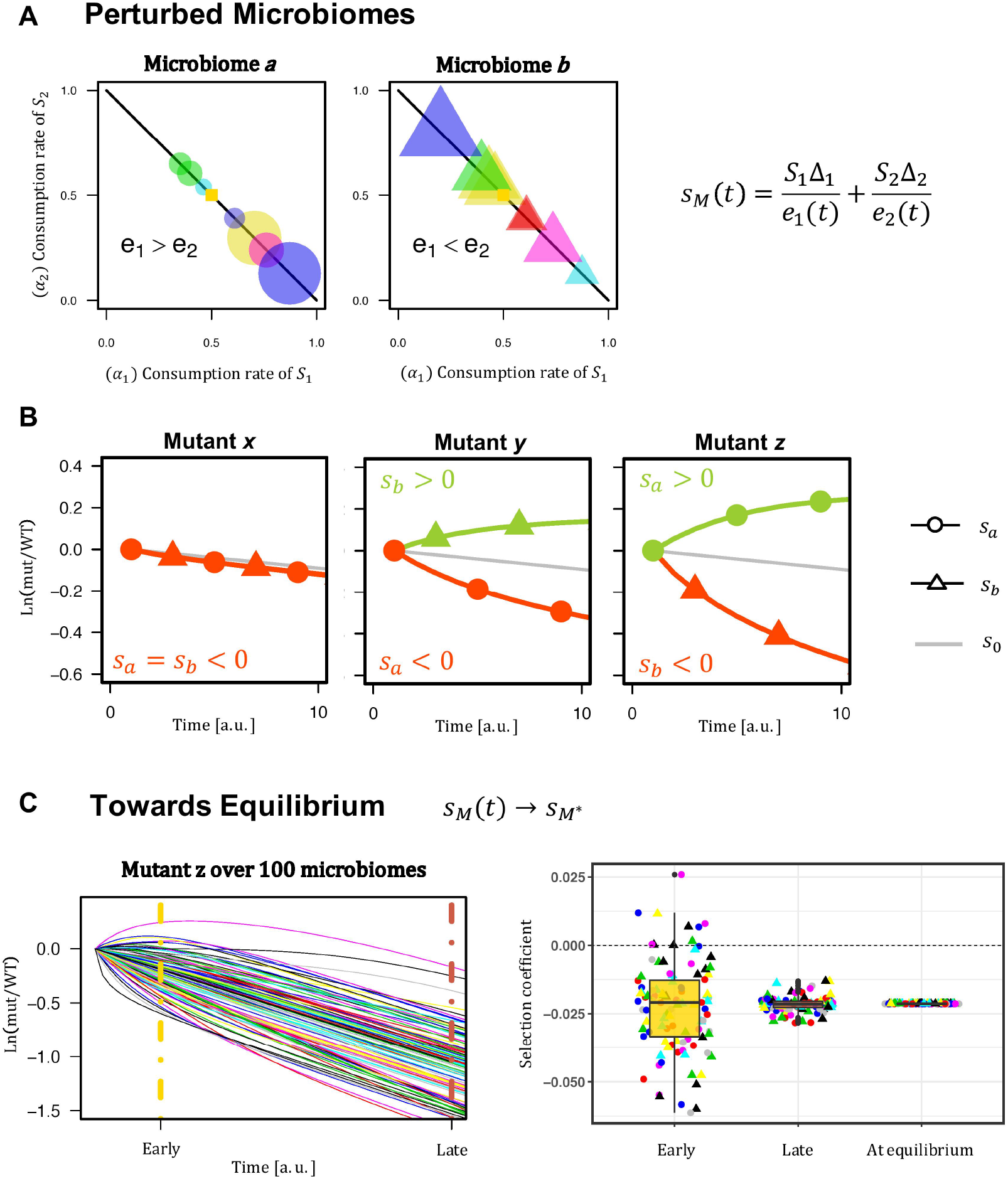
A general ecological model predicts time-dependent and host-specific selection on AR after antibiotic treatment. **(a)** Example of two microbiomes where a perturbation leads to functional distinct unbalances. Species in different colors with different relative abundances (represented as different areas of circles or triangles), at the colonization time; at equilibrium *e*_1_=*e*_2_ (Supplementary Text). **(b)** Selection depends on the mutation effect (mutant *x*,*y*, or *z*) and on the microbiome composition (*s_a_* or *s_b_*): for mutant *x*, which has same trait ratio as the WT, there is no time or microbiome dependence, whereas mutations *y* and *z* have opposite behaviors in the short time dynamics: selection is positive or negative depending on the microbial community. **(c)** Time-dependence of selection at short and long time-scales. As time passes and the microbiome moves towards equilibrium selection tends to a constant negative value (**Fig. 2c**): example of the cost dynamics of mutant *z* within 100 simulated microbiomes (equivalent dynamics but for mutants *x* and *y* in **Supplementary Fig. 6**).

### Compensatory evolution of AR strains

To experimentally test the theoretical prediction of time dependent compensatory evolution, we followed the long-term evolutionary dynamics of each AR clone colonizing the gut, after streptomycin treatment. Since the gut microbiota composition is more similar in mice from the same litter than mice from different litters^40,41^, our colonization experiment follows a design where the same AR background colonizes two mice from different parents (**Fig. 4A**). Thus, each mouse will likely differ in its microbiota composition state after antibiotic treatment is stopped (**Fig. 4b**). Analysis of the 16S rDNA in each colonized mouse indeed confirmed this expectation and significant differences between mice were found (**Fig. 4b**).

**Figure 4.**
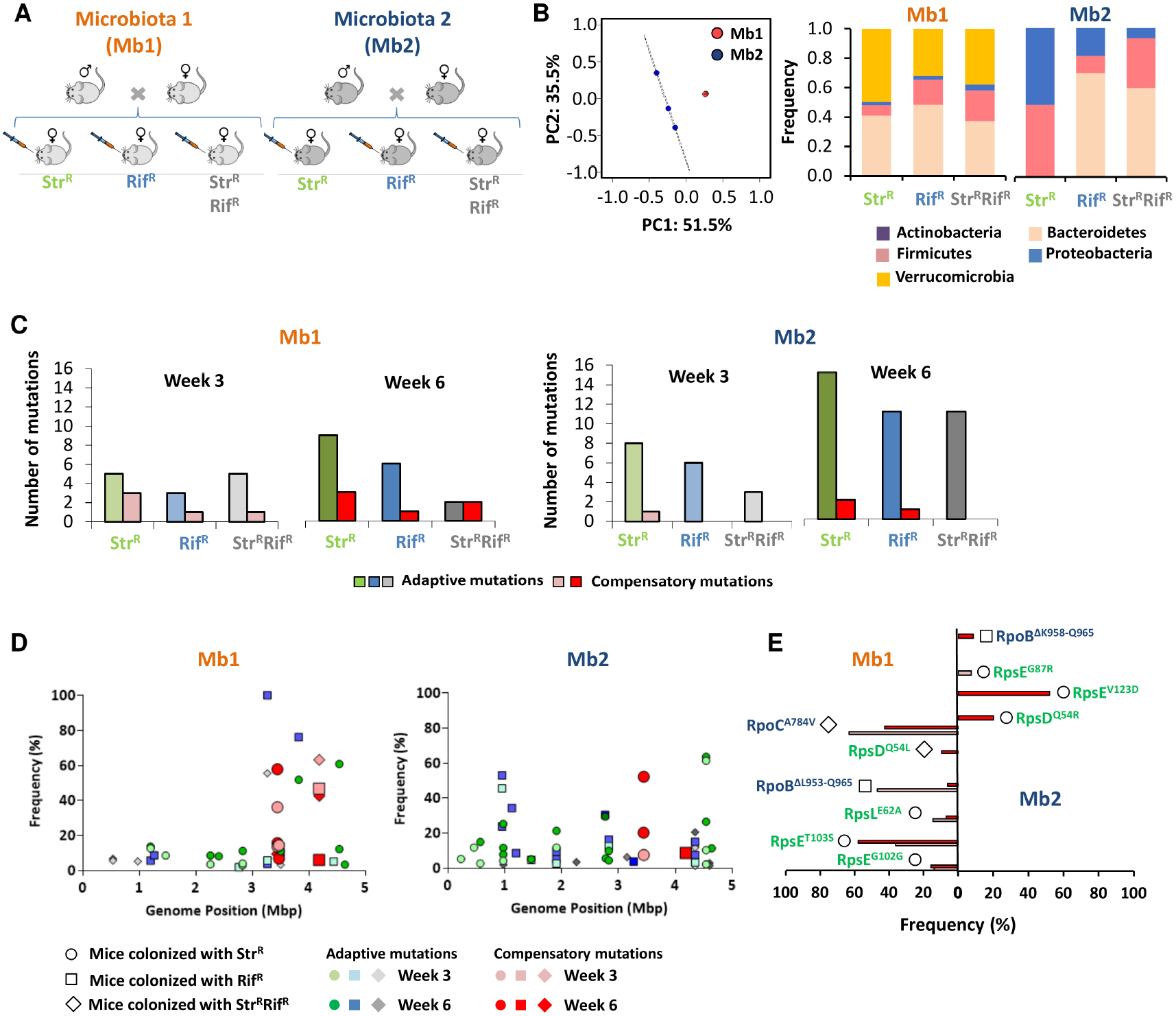
Dynamics and genetic basis of compensatory evolution of AR strains across hosts. **(a)** Experimental set up to study the adaption pattern of resistant strains (Str^R^, Rif^R^ and Str^R^Rif^R^) after an antibiotic perturbation. Mice from the same litter were co-housed for five weeks to homogenize the microbiota across litters. Afterwards, mice from the two different litters followed the same colonization resistance protocol as seen in Figure 1A and then used to follow adaptation of each of the resistant backgrounds (see Methods) for 6 weeks. Fecal samples were collected whole genome sequencing of populations and 16S. **(b)** Microbiota composition at the phylum level of the mice from the two different litters 3 weeks after colonization. Mice from the same litter cluster together and have a more similar microbiota. **(c)** Comparison of the number of putative adaptive and compensatory mutations present in the adapted resistant populations after 3 and 6 weeks in the mice gut with different microbiotas. **(d)** Frequency of the detected adaptive and compensatory mutations at week 3 and week 6. **(e)** Genetic basis of the bona fide compensatory mutations detected after 3 or 6 weeks of adaptation in the gut.

We next queried about the evolutionary dynamics of compensatory mutations along time and between hosts. To identify *bona fide* compensatory mutations we leveraged on the fact that these AR mutations have been extensively studied *in vitro*, in different media and bacterial species, and many of their targets have been identified^31,42–44^. Adaptive mutations unrelated to AR can also occur in the mouse gut at the time scale of weeks^35,45^ and many of those can be similar between mice with different microbiota compositions^37^. We thus expect adaptive mutations to be more similar across mice than compensatory mutations, which ought to be more specific to the AR background of the colonizing *E. coli*. Whole genome sequencing of pools of clones around week 3 and 6 after colonization reveal a temporal signal of compensatory evolution, and variation in the number of compensatory mutations between hosts. In the first cohort of mice at least one compensatory mutation could be detected in all AR backgrounds by the 3^rd^ week, whereas in the other cohort of mice no compensation for Rif^R^ or Srt^R^Rif^R^ could be detected at this earlier time point (**Fig. 4c**). This result is consistent with the expectation that after antibiotic perturbation, different microbiota compositions will reach equilibrium at different times and thus selection for the spread of compensatory mutations will be time dependent. Consistent with this interpretation, by the 6^th^ week the number of compensatory mutations detected increased in three out of the six studied mice. Remarkably no signal of compensatory evolution could be detected in one of the mice from cohort 2, even though 11 adaptive mutations raised in frequency in the double resistant lineage that colonized this host after six weeks of colonization (**Fig. 4d-e and Supplemental Table 5**). This data indicates that the cost of double resistance can take a long time to be expressed in specific hosts.

Analysis of the targets of evolutionary change and their frequency with the colonizing lineages showed that, in the majority of mice, adaptive mutations were more frequent than compensatory mutations, irrespective of the host or the AR genetic background (**Fig. 4d**). Overall 17 genes and 10 intergenic regions were called by natural selection for global adaptation across the 6 studied mice. Some of these have been shown to be adaptive when *E. coli* colonizes the gut of streptomycin-treated mice^35,45–47^.

The temporal pattern of population genomic variation strongly suggests that clonal interference between adaptive and compensatory mutations occurs. In some hosts the emerging compensatory mutations had weaker benefits than other adaptive mutations (e.g. mutations in *fimE* and *tdcA* reached higher frequencies than compensatory mutations to either Str^R^ or Rif^R^). The observed pattern is also consistent with the overall mutation rate to compensation being smaller than that of global adaptation to the gut. rpoB and rpoC were the two targets for compensation to Rif^R^, with deletions in rpoB being alleles that have not been commonly detected *in vitro*. The three targets for compensation to Str^R^: in the *rpsE*, *rpsL* and *rpsD* genes have been detected in previous studies of compensation under laboratory conditions (**Fig. 4d-e** and **Supplementary Table 5**).

Overall these observations are consistent with the observed variation of fitness effects of AR in the short-term competitions (**Fig. 1b**) and with the results of the simple theoretical model described above (**Fig. 2–3**), predicting a strong personalized pattern of compensation for deleterious pleiotropic mutations at the initial stages of evolution (**Fig. 4**).

## DISCUSSION

Many chromosomal encoded AR exhibit a fitness cost when growing in *in vitro* artificial laboratory environments. How previously measured costs of AR *in vitro* translate into the natural environments is currently poorly known. Yet, the quantification of the strength of selection for and against resistance in ecosystems such as the mammalian gut is critical for understanding the levels of the microbiota resistome^8^. In the species rich intestinal tract, bacteria ferociously compete for resources and the environment may not be as constant as that of laboratory settings. Indeed, we have found that the costs of both single and double resistances *in vitro* could significantly deviate from their estimated *in vivo* effects, even in the simplest case of mono-colonized hosts (**Fig. 1b** and **Supplementary Fig. 3**). In the more relevant model of *E. coli* colonization of a complex gut microbiota with inter-species interactions, we uncovered that the same AR mutation can have a wide range of fitness effects in hosts that are genetically identical, eat the same diet and experience the same environment. Following antibiotic treatment, a given AR mutation showed a strong deleterious effect in competitive fitness within one host but increased in frequency in another - a display of antagonistic pleiotropy. A similar finding occurs when double resistant strains compete with single resistant lineages (**Supplementary Figure 2**). Since the host specific effect is absent in germ-free mice (**Fig. 1c**), our observations suggest that selection against resistance is acting unequally across mice due to the presence of the microbiota. In accordance with previous reports^48,49^, a decrease in microbiota diversity following antibiotic treatment is also seen in our study, as well as a high variance of its composition between mice (**Fig. 1e-f**). Differences in the composition of the microbiota can lead to differences in metabolic activity of the whole ecosystem, which in turn will likely result in distinct levels of inter-species competition for the different gut resources. Since the AR studied here, involving changes in the ribosome and RNA polymerase, affect metabolism^50–52^, it is natural to expect that their fitness effects may depend on the microbiota composition, as observed. Streptomycin resistance mutations can affect translation speed and accuracy^53^, while certain *rpoB* mutations can affect transcription speed^54^ and fidelity^55,56^. Cellular processes that depend of the effectiveness of transcription and translation, such as the activation or repression of operons linked to nutrient uptake and consumption, are likely to be affected, generating distinct consumption rates when compared with the wild-type strain. Accordingly, the mutations under study have been shown to change the growth and competitive fitness of bacteria in different nutritional environments^22^, suggesting that they can change the relative consumption of different resources in natural environments. By theoretical modeling the effect of AR mutations in a framework of competition within ecosystems, we found that AR mutations, which are costly in the absence of interspecies interactions, should entail variable costs in the context of host-specific microbiota perturbations. The model also predicted time dependent-selection of the fate of AR, implying that the strength of selection to lower resistance costs should generally show variation along time within and between individuals. Such patterns were corroborated by *in vivo* long-term evolution experiments on three different resistance backgrounds. Notwithstanding other key simplifications in the model, we also did not explicitly consider the emergence of compensatory mutations nor the clonal interference pattern observed during the long-term evolutionary dynamics of *E. coli* resistant strains in the gut. The quantitative understanding of how globally adaptive mutations interfere with background specific compensatory mutations during the eco-evolutionary dynamics of gut commensal bacteria after microbiota dysbiosis is an important problem for future theoretical work, that can illuminate specific *in vivo* experimental evolution designs.

The findings that metabolic adaptations occur in every host, that typical compensatory mutations may take a long time to reach high frequency, and that reversion to a sensitive state are not detected, has consequences for the expansion and maintenance of resistant strains within hosts. A recent study showing that a short-term cefuroxime treatment can increase the general level of resistance in the human gut microbiota^57^, corroborates this expectation, although the factorial level of potential causes for such an effect is enormous when studying AR levels in humans.

The dysbiotic period following antibiotic treatment offers a time window of opportunity for disease causing bacteria to invade the host intestine. The associated possible reduction in costs of resistance at this critical period offers an important breach for the maintenance of resistant pathogens, and further difficulties in elimination of these agents. Yet in the case of AR mutations that affect nutritional metabolism, an interesting possibility of using specific dietary supplementation could be considered. As metabolic model predictions from genomic data of strains is rapidly improving and specific carbon supplementations can sometimes be effective in changing the frequency of specific strains^58^, hopes that such progress can be harnessed to lower resistance levels may become within reach.

A study with a simplified model microbiota has shown that the presence of a single gut bacterial species can change the outcome of intra-species competitions^59^. Therefore, a plausible strategy to eliminate resistant pathogens is to find competitors that will reliably and specifically generate a cost for the resistant strain. Studying the fitness effects of resistance mutations in the presence of specific gut microbes or defined collections of microbiota members, and further testing the efficiency of these strains in dysbiosis models could lead to optimized approaches for selection against resistance.

According to our model, multi-species can coexist when there are, at least, two resources available for which the different species compete. Importantly, the species are able to consume both resources, even though they have different abilities to consume each one. There is evidence that several gut species can use more than one carbon sugar^60^. Even though carbon-catabolite-repression (CCR) is known to occur in *E. coli* for carbon sources such as glucose^61^, bacteria can find a multitude of nutritional niches when colonizing the mammalian gut. Successful species probably evolved to be versatile enough to switch their realized nutrient niche regularly or to simultaneously utilize multiple substrates^62^. In agreement, the gene expression profiles *E. coli* MG1655 grown in mucus (mimicking the gut nutritionally) identified genes involved in catabolism of several sugars such as N-acetylglucosamine, sialic acid, glucosamine, fucose, ribose, glucuronate, galacturonate, gluconate, and maltose^63^.

We found evidence for significant antagonistic pleiotropy for AR fitness effects between hosts. Antagonistic pleiotropy could also occur within a single host intestine, as individual *E. coli* cells might experience different niches in such structured environment, while the population as a whole may consume different carbon sources simultaneously^58^. The simple theoretical model of resource competition helps explaining how the host-dependent AR costs can arise from general properties of the ecosystem even without species-specific or cross-feeding interactions. Pairwise cross-feeding interactions between gut bacteria can nevertheless occur^64,65^ and higher order cross-feeding interactions are thought to be involved in complex microbial communities^66^. While cross-feeding interactions are a feature of the gut ecosystem, these networks of metabolites produced by bacteria can also be affected by strong microbiota perturbations. Thus, an altered ability to consume cross-fed resources by resistant bacteria would lead to the same outcome: a host-specific fitness effect of resistance mutations in dysbiosis.

Recent studies that define the bacterial taxa within human microbiota demonstrate significant variability between individuals^67,68^. One of these studies^68^ was able to track individuals from hundreds of people by using the microbiota data available in the “Human Microbiome Project” database. This is strong evidence that our microbiota has enough unique characteristics to be almost used as a “fingerprint” of an individual. As the microbiota can affect the cost of resistance, it is likely that the fate of resistant bacteria in humans is also host-specific. Therefore, depending on individual microbiomes and resistomes, the fight against antibiotic resistance in the current era might require personalized medicine.

## METHODS

### *E. coli* and mice strains

All of the strains were derived from *Escherichia coli* strain K-12 MG1655. Since the gat operon was observed to be a mutation hotspot under strong selection for our strains in the mouse gut ^35,45^, we pre-adapted our *E. coli* strain to a *gat* negative phenotype by knocking down the *gatZ* gene permanently. Briefly, P1 transductions were performed in order to delete the *gatZ* gene from our strains as a pre-adaptive mutation and strain *E. coli* JW2082-1 from the KEIO collection was used as a donor.

The new strains, LC88 and RB929 (*ΔlacIZYA::scar galK::cat-YFP/CFP ΔgatZ::FRT-aph-FRT*), were used as wild-type strains in the competitions. P1 transduction was also used to insert the point mutation *rpoB*^H526Y^ (Rif^R^) in the wild-type background and to pass the *gatZ* deletion from the wild-type strains to isogenic antibiotic resistant strains, carrying either the point mutation *rpsL*^K43T^ (Str^R^) or both *rpsL*^K43T^ and *rpoB*^H526Y^ mutations (Str^R^Rif^R^). The resulting streptomycin resistant (Str^R^) strains LC81 and LC82 (YFP/CFP, respectively), the rifampicin resistant (Rif^R^) strains RB933 and LC84b (YFP/CFP, respectively), and the double resistant (Str^R^Rif^R^) strains LC85, LC86 (YFP/CFP, respectively) were used to colonize the mice and perform the competitions *in vivo*.

Six-to-thirteen week-old female C57BL/6J germ-free (GF) mice were used as hosts for the *in vivo* competitions in the absence of microbiota, while 6-to-8 week-old female C57BL/6J specific pathogen free (SPF) mice were used for the *in vivo* competitions and the evolution experiment in the presence of microbiota. GF mice were bred and raised at the IGC gnotobiology facility in dedicated axenic isolators (La Calhene/ORM).) Young adults were transferred into sterile ISOcages (Tecniplast) before the competition experiments.

### *In vitro* competitions

The strains were streaked from the frozen stocks into LB agar with antibiotics corresponding to their resistance and incubated at 37°C for 24 hours, followed by acclimatization for 24h in LB and in minimal media with 0.4% glucose, in 96-well plates, at 37°C, with shaking (700 rpm). Each resistant strain was mixed in a 1:1 ratio with the sensitive wild-type, and competitions were performed for 24h in the same conditions as the acclimatization. To determine the initial and final ratios of resistant and susceptible strains in the competition assays, bacteria were quantified with an LSR Fortessa flow cytometer using a 96-well plate autosampler. Samples were always run in the presence of SPHERO (AccuCount 2.0-μm blank particles) in order to accurately quantify bacterial numbers in the cultures. Briefly, flow cytometry samples consisted of 180 μl of PBS, 10 μl of SPHERO beads, and 10 μl of a 100-fold dilution of the bacterial culture in PBS. The bacterial concentration was calculated based on the known number of beads added. Cyan fluorescent protein (CFP) was excited with a 442-nm laser and measured with a 470/20-nm pass filter. Yellow fluorescent protein (YFP) was excited using a 488-nm laser and measured using a 530/30-nm pass filter. The selection coefficient (s) of each mutant strain was estimated as the per generation (number of doublings of the susceptible strain) difference in the ration of the resistant strain and the reference strain after 24h: S = ln(R_f_/R_i_)/t, where t corresponds to the number of generations and R_f_ and R_i_ to the final and initial ratios between resistant and reference strains, respectively. The gat negative phenotype had no interference in between the negative epistasis in between resistances ^9,22^.

### *In vivo* competitions

To measure the fitness effects and to evolve the resistant strains in SPF mice, we used a streptomycin treatment in order to break the colonization resistance. Mice were separated into individual cages and given autoclaved drinking water containing streptomycin sulfate (5g/L) for seven days and then mice were given regular autoclaved drinking water for 2 days, in order to wash out the antibiotic from the gut and allow for the microbiota stabilization. After 4 hours of starvation for food and water, the mice were gavaged with 100 μl of a ≈10^9^ cells/ml suspension with a 1:1 ratio of the two competing strains.

To make the suspension, the strains were streaked from stocks in LB agar with antibiotics corresponding to their resistance two days before gavage and incubated for 24 hours, followed by an overnight culture of a single colony for each biological replicate in BHI (brain heart infusion) media with the corresponding antibiotic. The cultures were then diluted 100-fold and grown in BHI media until an OD_600nm_ ≈ 2. Flow cytometry was used to assess the number of cells per growth and therefore adjust the initial number of cells in order to prepare the suspension for the gavage. The same protocol was used in order to generate the bacteria suspension for the GF mice. Mice fecal pellets were collected 4 hours and every 24 after gavage, for 5 days, suspended and diluted in PBS and plated in LB agar plates. Plates were incubated overnight and the frequencies of CFP- or YFP-labeled bacteria were assessed by counting the fluorescent colonies with the help of a fluorescent stereoscope (SteREOLumar, Carl Zeiss). The samples were also stored in 15% glycerol at −80°C for future experiments. The selection coefficient (S) per day of each mutant strain was estimated through the slope of the log-linear regression of the ratio of the resistant strain and the reference strain from day 1 to day 5. Apart from the streptomycin treatment to break colonization resistance, the same protocol was used in the competitions with GF mice.

### Microbiota analysis

To assess the gut microbiota composition of mice, we extracted DNA from fecal samples from two experiments: the measurement of fitness costs in SPF mice (**Fig. 1**) and from the compensatory evolution of the resistant strains (**Fig. 4**). For the analysis of the microbiota perturbation during the measurement of the fitness costs, fecal samples were collected from 8 mice belonging to different litters, before the start of the antibiotic treatment and 72 hours after its end, corresponding to the first time-point on the competition experiments. For samples collected during the *E. coli* resistant strains compensatory evolution experiments, we characterized the microbiota context by analyzing the fecal samples collected from 6 female mice from two different litters (3 per litter) at day 17 of the evolution experiment.

Fecal DNA was extracted with a QIAamp DNA Stool MiniKit (Qiagen), according to the manufacturer’s instructions and with an additional step of mechanical disruption^69^. 16S rRNA gene amplification and sequencing was carried out at the Gene Expression Unit from Instituto Gulbenkian de Ciência, following the service protocol. For each sample, the V4 region of the 16 S rRNA gene was amplified in triplicate, using the primer pair F515/R806, under the following PCR cycling conditions: 94 °C for 3 min, 35 cycles of 94 °C for 60 s, 50 °C for 60 s, and 72 °C for 105 s, with an extension step of 72 °C for 10 min. Samples were then pair-end sequenced on an Illumina MiSeq Benchtop Sequencer, following Illumina recommendations.

QIIME2^70^ was used to analyze the 16S rRNA sequences by following the authors’ online tutorials (https://docs.qiime2.org/2018.11/tutorials/). Briefly, the demultiplexed sequences were filtered using the “denoise-single” command of DADA2^71^, and forward and reverse sequences were trimmed in the position in which the 25^th^ percentile’s quality score got below 20. Alpha diversity and ANCOM analysis^72^ were performed as in the QIIME2 tutorial. Beta diversity distances were calculated through Unweighted Unifrac^73^, and PCoA on the respective distance matrices were performed using the R software ((http://www.R-project.org) and the R packages “vegan” (https://CRAN.R-project.org/package=vegan), “BiodiversityR” (https://CRAN.R-project.org/package=BiodiversityR) and “RVAideMemoire” (https://CRAN.R-project.org/package=RVAideMemoire). For taxonomic analysis, OTU were picked by assigning operational taxonomic units at 97% similarity against the Greengenes database^74^.

### Compensatory evolution in SPF mice

To study the adaptation of resistance strains to the gut, three sister mice from 2 different litters were used, for a total of 6 mice. For each resistant genotype, we colonized 1 mouse from each litter with a mix of YFP and CFP-labeled bacteria. The whole colonization protocol was identical to the *in vivo* competitions as described for the SPF mice. Samples were collected 24h after gavage and every 48h thereafter, until 39 days post colonization. All samples were stored in 15% glycerol at −80°C.

### DNA extractions and whole-genome sequencing analysis

Concentration and purity of DNA were quantified using Qubit and NanoDrop, respectively. The DNA library construction and sequencing was carried out by the IGC genomics facility. Each sample was pair-end sequenced on an Illumina MiSeq Benchtop Sequencer. Standard procedures produced data sets of Illumina paired-end 250 bp read pairs. The reads were filtered using SeqTk version 1.0-r63. The mean coverage after filtering for the different samples was as follows: 168x and 175x for Str^R^1 day 19 and day 39, respectively; 238x and 194x for Str^R^2 day 19 and day 39, respectively; 164x and 159x for Rif^R^1 day 19 and day 39, respectively; 226x and 202x for Rif^R^2 day 19 and day 39, respectively; 148x and 156x for Str^R^ Rif^R^1 day 19 and day 39, respectively; 213x and 220 for Str^R^ Rif^R^2 day 19 and day 39, respectively. Sequences were analyzed using Breseq version 0.31.1, using *E. coli* K12 genome NC_000913.3 as a reference, with the polymorphism option selected, and the following parameters: (a) rejection of polymorphisms in homopolymers of a length greater than three, (b) rejection of polymorphisms that are not present in at least three reads in each strand, and (c) rejection of polymorphisms that do not have a p-value for quality greater than 0.05,(d) rejection of polymorphisms with less than 3 of coverage in each strand and (e) rejection of polymorphisms with less than 1% frequency. All other Breseq parameters were used as default. Hits that were present in all of our ancestral mutants as well as homopolymers were discarded. Hits that were likely to be due to misalignment of repetitive regions were also discarded. Regarding the downstream analysis, target genes that appeared only in one sample and had a frequency lower than 5% were not considered.

### Modeled AR competitions

Numerical simulations were used to confirm the analytical predictions and to graphically represent the results. The dynamics of M species competing for P resources follow a recent formalization of the classical MacArthur consumer-resource model^39^:

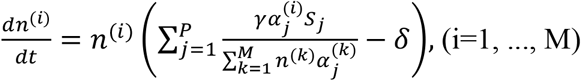

Where *n*^(*i*)^ (*t*) is the density of species *i*,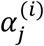 is the consumption rate of substrate *j* by species *i*, *S*_*j*_ is the constant substrate *j* supply, *γ* is the yield and *δ* is the microbial death rate. A detailed description of the parameter choice and the algorithm is given in the supplementary information. The implementation of the competitions and the graphical resolution of the two-resource scenario were done in RStudio 1.1.463 and the source code is available upon request to the authors.

### Statistical analysis

The selection coefficient of the *in vivo* competitions was tested in R software, through an F-statistic on a predictive linear model of the mutant/sensitive or double mutant/single mutant ratio over time, generated through the observed ratio on sampled time-points from 24, 48, 72, 96 and 120 hours after gavage. The null hypothesis is that the slope, which is an estimation of the selection coefficient, is equal to 0. When the null hypothesis was rejected, p-value < 0.05, the mutant was considered to have a cost if the slope of the model was negative and to have a fitness benefit if the slope was positive. F tests were performed to analyze the variance in between hosts. Normality of each treatment was tested through with Shapiro Wilk test and normality of the treatments involving competitions in the presence of microbiota was further tested through Kolmogorov-Smirnov test.

### Ethics statement

This research project was ethically reviewed and approved by the Ethics Committee of the Instituto Gulbenkian de Ciência (license reference: A009.2018) and by the Portuguese National Entity that regulates the use of laboratory animals (DGAV – Direção Geral de Alimentação e Veterinária (license reference: 008958). All experiments conducted on animals followed the Portuguese (Decreto-Lei n° 113/2013) and European (Directive 2010/63/EU) legislations, concerning housing, husbandry and animal welfare.

## ACKNOWLEDGMENTS

LLC, PD and MA were supported by “Fundação para a Ciência e Tecnologia” (FCT), fellowships SFRH/BPD/118474/2016, PD/BD/106003/2014 and PD/BD/138735/2018, respectively. Research was supported by project JPIAMR/0001/2016-ERA NET and ONEIDA project (LISBOA-01-0145-FEDER-016417) co-funded by FEEI – “Fundos Europeus Estruturais e de Investimento” from “Programa Operacional Regional Lisboa 2020”, and by national funds from FCT. We would like to thank the personnel of the IGC Rodent Facility, Genomic Facility and the Bioinformatics Unit for their assistance.

## AUTHOR CONTRIBUTIONS

LLC and PD performed the experiments. MA and IG designed the model which was developed by MA. LLC, PD and IG analyzed the results. IG coordinated the study. All authors contributed in the writing of the manuscript and gave final approval for publication.

