## SupplementalText for "Dysbiosis personalizes fitness effect of antibiotic resistance in the mammalian gut"

### SUPPLEMENTARY TABLES

**Supplementary Table 1 – Selective effect estimated from the competitions of AR in specific pathogen free (SPF) mice.** Estimates were obtained from the slope of ln[Resistant /Susceptible] across 5 days of co-colonization in each mouse. Fitness effects with a significant positive (negative) slope indicate that the resistance had a beneficial (deleterious) effect over this time period. Significant slopes are indicated as *. Fitness effects were converted to generations assuming generation time of *E. coli* colonizing specific-pathogen-free mice[1].

| Mouse | Competitors | $S_{day}$ (2.SE) | $S_{gen}$ (2.SE) | $R^2$ |
|---|---|---|---|---|
| S1 | **$Str^R$ vs Wt** | -1.4 (0.4)* | -0.08 (0.02)* | 0.93 |
| S2 | **$Str^R$ vs Wt** | 0.8 (0.4)* | 0.04 (0.02)* | 0.25 |
| S3 | **$Str^R$ vs Wt** | -1.3 (0.2)* | -0.07 (0.02)* | 0.77 |
| S4 | **$Str^R$ vs Wt** | 0.07 (0.16) | 0.004 (0.008) | 0.24 |
| S5 | **$Str^R$ vs Wt** | -0.7 (0.2)* | -0.04 (0.01)* | 0.79 |
| S6 | **$Str^R$ vs Wt** | -0.2 (0.1)* | -0.012 (0.006)* | 0.90 |
| R1 | **$Rif^R$ vs Wt** | -0.71 (0.08)* | -0.037 (0.004)* | 0.99 |
| R2 | **$Rif^R$ vs Wt** | 0.2 (1.2) | 0.01 (0.07) | 0.03 |
| R3 | **$Rif^R$ vs Wt** | -0.4 (0.4) | -0.02 (0.04) | 0.58 |
| R4 | **$Rif^R$ vs Wt** | -0.2 (0.2) | -0.01 (0.01) | 0.62 |
| R5 | **$Rif^R$ vs Wt** | 0.5 (0.6) | 0.03 (0.03) | 0.51 |
| R6 | **$Rif^R$ vs Wt** | -1.1 (0.4)* | -0.06 (0.02)* | 0.90 |
| SR1 | **$Str^R$ $Rif^R$ vs Wt** | -0.8 (0.2)* | -0.04 (0.01)* | 0.94 |
| SR2 | **$Str^R$ $Rif^R$ vs Wt** | -0.6 (0.6) | -0.03 (0.03) | 0.58 |
| SR3 | **$Str^R$ $Rif^R$ vs Wt** | -0.4 (0.4) | -0.02 (0.02) | 0.70 |
| SR4 | **$Str^R$ $Rif^R$ vs Wt** | 0.4 (0.4) | 0.02 (0.02) | 0.60 |
| SR5 | **$Str^R$ $Rif^R$ vs Wt** | -0.3 (0.2) | -0.01 (0.01) | 0.55 |
| SR6 | **$Str^R$ $Rif^R$ vs Wt** | -0.4 (0.6) | -0.02 (0.3) | 0.39 |
| DS1 | **$Str^R$ $Rif^R$ vs $Str^R$** | -0.39 (0.07)* | -0.02 (0.004)* | 0.98 |
| DS2 | **$Str^R$ $Rif^R$ vs $Str^R$** | -0.03 (0.08) | -0.002 (0.002) | 0.17 |
| DS3 | **$Str^R$ $Rif^R$ vs $Str^R$** | -0.4 (0.4) | -0.02 (0.02) | 0.61 |
| DR1 | **$Str^R$ $Rif^R$ vs $Rif^R$** | 0 (0.2) | 0 (0.01) | 0.00 |
| DR2 | **$Str^R$ $Rif^R$ vs $Rif^R$** | 0.3 (0.1)* | 0.015 (0.004)* | 0.93 |
| DR3 | **$Str^R$ $Rif^R$ vs $Rif^R$** | -1.3 (0.2)* | -0.07 (0.01)* | 0.98 |

**Supplementary Table 2 – Temporal sampling to test for phenotypic reversion during the evolution experiment.** Stool samples were diluted and grown in LB with chloramphenicol to select *E. coli* strains. Random colonies were replicated into LB plates with streptomycin, with rifampicin, with both drugs and with no antibiotics. No reversions were observed in the tested clones.

| Population/litter | Day 11 | Day 17 | Day 19 | Day 25 | Day 29 | Day 39 | Total | Reversions |
|---|---|---|---|---|---|---|---|---|
| **$Str^R$ 1** | 102 | 153 | | 152 | 113 | 115 | **635** | **0** |
| **$Rif^R$ 1** | 110 | 100 | | 96 | 96 | 96 | **498** | **0** |
| **$Str^RRif^R$ 1** | 78 | 135 | | 132 | 114 | 102 | **561** | **0** |
| **$Str^R$ 2** | | | 144 | | | | **144** | **0** |
| **$Rif^R$ 2** | | | 96 | | | | **96** | **0** |
| **$Str^RRif^R$ 2** | | | 144 | | | | **144** | **0** |

**Supplementary Table 3 – Fitness effects of resistance mutations in germ-free (GF) mice.** Fitness effects of resistances - $Str^R$ (*rpsL*$^{K43T}$), $Rif^R$ (*rpoB*$^{H526Y}$) and $Str^R Rif^R$ (*rpsL*$^{K43T}$*rpoB*$^{H526Y}$) - against a susceptible strain and of the double resistant against the single resistant mutants. Errors represent 2*SE. Epistasis was measured using the additive model. The error for epistasis was calculated through the error propagation of the standard errors.

| Competition | $S_{/gen}$* Germ-Free | Epistasis |
|---|---|---|
| **$Str^R$ vs Wt** | -0.008 ± 0.003 | |
| **$Rif^R$ vs Wt** | -0.009 ± 0.003 | |
| **$Str^R Rif^R$ vs $Str^R$** | -0.010 ± 0.003 | |
| **$Str^R Rif^R$ vs $Rif^R$** | -0.008 ± 0.002 | |
| **$Str^R Rif^R$ vs Wt** | -0.017 ± 0.004 | 0.000 ± 0.003 |

*assuming the generation time of *E. coli* colonizing germ-free mice measured previously[1].

**Supplementary Table 4 – Selective effect estimated from the competitions of AR in germ-free mice.** Estimates were obtained from the slope of ln[Resistant /Susceptible] across 5 days of co-colonization in each mouse. Fitness effects with a significant negative slope indicate that the resistance had a deleterious effect over this time period. Conversion of fitness effects per day to generations was done assuming generation time of *E. coli* colonizing germ-free mice[1]. All resistances have a significant cost across all mice.

| Mouse | Competitors | $S_{day}$ (2.SE) | $S_{day}$ (2.SE) | $R^2$ |
|---|---|---|---|---|
| S1 | **$Str^R$ vs Wt** | -0.17 (0.08) | -0.007 (0.004) | 0.84 |
| S2 | **$Str^R$ vs Wt** | -0.3 (0.1) | -0.011 (0.004) | 0.92 |
| S3 | **$Str^R$ vs Wt** | -0.14 (0.06) | -0.005 (0.002) | 0.87 |
| R1 | **$Rif^R$ vs Wt** | -0.28 (0.08) | -0.011 (0.002) | 0.94 |
| R2 | **$Rif^R$ vs Wt** | -0.2 (0.1) | -0.006 (0.004) | 0.78 |
| R3 | **$Rif^R$ vs Wt** | -0.30 (0.06) | -0.011 (0.002) | 0.98 |
| SR1 | **$Str^R$ $Rif^R$ vs Wt** | -0.39 (0.04) | -0.015 (0.02) | 0.99 |
| SR2 | **$Str^R$ $Rif^R$ vs Wt** | -0.5 (0.1) | -0.021 (0.006) | 0.95 |
| SR3 | **$Str^R$ $Rif^R$ vs Wt** | -0.38 (0.06) | -0.015 (0.002) | 0.98 |
| DS1 | **$Str^R$ $Rif^R$ vs $Str^R$** | -0.32 (0.08) | -0.012 (0.002) | 0.97 |
| DS2 | **$Str^R$ $Rif^R$ vs $Str^R$** | -0.2 (0.1) | -0.008 (0.004) | 0.83 |
| DS3 | **$Str^R$ $Rif^R$ vs $Str^R$** | -0.26 (0.06) | -0.010 (0.001) | 0.95 |
| DR1 | **$Str^R$ $Rif^R$ vs $Rif^R$** | -0.18 (0.04) | -0.007 (0.002) | 0.96 |
| DR2 | **$Str^R$ $Rif^R$ vs $Rif^R$** | -0.26 (0.02) | -0.010 (0.001) | 0.99 |
| DR3 | **$Str^R$ $Rif^R$ vs $Rif^R$** | -0.19 (0.04) | -0.007 (0.002) | 0.98 |

2.SE – 2 x standard error.

**Supplementary Table 5 – *De novo* mutations on AR *E. coli* genetic backgrounds during gut colonization.** Genome position, mutation target and mutation frequency after 3 and 6 weeks of evolution are shown. Mutations in bold occurred in genes identified in previous literature as targets for compensation. References on mutational gene targets previously described in the literature are indicated after the mutation target.

| | | | | Frequency | |
|---|---|---|---|---|---|
| **Population** | **Genome Position** | **Gene** | **Mutation** | **Week 3** | **Week 6** |
| **$Str^R$ 1** | 1198436 | ymfE / lit | +AATGAAAT | 12.6% | 13.6% |
| | 1466201 | ydbA[2] | T→C | 8.5% | |
| | 2259422 | psuK / fruA[3] | A→C | 3.5% | 8.5% |
| | 2406600 | lrhA | +TCGAGG | | 8.1% |
| | 2829125 | srlR[4] | C→T | 2.9% | |
| | 2829468 | srlR | C→A | 3.9% | 11.1% |
| | **3444923** | **rpsE**[5,6] | **T→G** | **13.7%** | **15.6%** |
| | **3444925** | **rpsE**[5,6] | **C→G** | **36.1%** | **57.8%** |
| | **3474368** | **rpsL**[5,6] | **T→G** | **14.4%** | **6.8%** |
| | 3504197 | frlR[4] | G→A | | 10.9% |
| | 3823019 | spoT[2] | G→A | | 51.8% |
| | 4542161 | fimE[3] | IS5 +4 bp | | 60.8% |
| | 4542457 | fimE[3] | IS1 +9 bp | | 12.2% |
| | 4640748 | yjjY → / → yjtD[7] | IS5 +4 bp | | 3.3% |
| **$Str^R$ 2** | 228767 | rrfH | C→A | 5.2% | |
| | 457912 | clpX | IS1 +9 bp | 11.7% | |
| | 570603 | ybcK → / → ybcL | IS2 +5 bp | 2.6% | 14.9% |
| | 972965 | elyC | G→T | | 11.5% |
| | 973071 | elyC | IS1 +9 bp | | 25.2% |
| | 973103 | elyC | IS5 +4 bp | 4.1% | 25.1% |
| | 973108 | elyC | G→T | | 7.7% |
| | 1466210 | ydbA[2] | G→A | | 5.0% |
| | 1466438 | ydbA[2] | T→G | | 5.0% |
| | 1909535 | kdgR[4] | G→A | 11.7% | 21.3% |
| | 2765463 | yfjL ← / ← yfjM | +TATGGCAC | | 29.4% |
| | 2773761 | yfjW | T→C | | 5.5% |
| | 2829065 | srlR[4] | C→T | | 9.8% |
| | 2829207 | srlR[4] | G→A | | 4.4% |
| | **3441515** | **rpsD**[5,6] | **T→C** | | **20.2%** |
| | **3444862** | **rpsE**[5,6] | **A→G** | | **52.1%** |
| | **3444971** | **rpsE**[5,6] | **C→T** | **7.5%** | |
| | 4542308 | fimE[3] | IS5 +4 bp | 2.1% | 26.5% |
| | 4542308 | fimE[3] | IS5 +4 bp | 10.2% | |
| | 4542577 | fimE[3] | IS1 +10 bp | 61.3% | 63.5% |
| | 4640605 | yjjY → / → yjtD[7] | IS2 +5 bp | | 11.3% |

| | | | | | |
|---|---|---|---|---|---|
| **Rif$^R$ 1** | 1198437 | ymfE ← / → lit | Δ8 bp | | 5.6% |
| | 1264717 | hemA | G→T | | 8.6% |
| | 2765412 | yfjL ← / ← yfjM | Δ8 bp | 1.8% | |
| | 3266935 | tdcA | T→C | 5.7% | |
| | 3266993 | tdcA | C→T | | 100.0% |
| | 3267147 | tdcA ← / → tdcR[4] | IS5 +4 bp | | 3.8% |
| | 3504139 | frlR[4] | C→T | | 10.4% |
| | 3822706 | spoT[2] | C→T | | 76.1% |
| | **4184101** | **rpoB**[8,9] | **Δ39 bp** | **46.7%** | **6.0%** |
| | 4439596 | ytfK | T→A | 5.0% | |
| **Rif$^R$ 2** | 954669 | focA ← / ← ycaO[4] | IS5 +4 bp | 45.5% | 52.9% |
| | 954678 | focA ← / ← ycaO[4] | IS5 +4 bp | | 23.7% |
| | 1125778 | rimJ[4] | IS1 +9 bp | | 34.3% |
| | 1198460 | ymfE ← / → lit | +AATGAAAT | | 8.5% |
| | 1466432 | ydbA[2] | A→G | 4.6% | |
| | 1466438 | ydbA[2] | T→G | 4.7% | |
| | 1909925 | kdgR[4] | IS5 +4 bp | | 5.0% |
| | 1910031 | kdgR[4] | IS2 +5 bp | 2.5% | 9.4% |
| | 2765411 | yfjL ← / ← yfjM | +GCACTATG | | 30.4% |
| | 2829687 | srlR[4] | A→C | 12.4% | 16.2% |
| | 3267147 | tdcA ← / → tdcR[4] | IS5 +4 bp | | 3.8% |
| | **4184117** | **rpoB**[8,9] | **Δ24 bp** | | **8.6%** |
| | 4348862 | dcuB ← / ← dcuR[4] | IS5 +4 bp | 3.2% | 7.4% |
| | 4349082 | dcuB ← / ← dcuR[4] | IS2 +5 bp | | 15.0% |
| **Str$^R$ Rif$^R$ 1** | 533245 | allR | T→G | 5.6% | 6.7% |
| | 974859 | mukF | Δ3 bp | 5.2% | |
| | 3267149 | tdcA ← / → tdcR[4] | Δ2 bp | 55.5% | |
| | **3441515** | **rpsD**[5,6] | **T→A** | | **9.4%** |
| | 3504101 | frlR[4] | G→T | 15.2% | 8.2% |
| | 3504565 | frlR[4] | C→T | 3.2% | |
| | **4187700** | **rpoC**[8,9] | **C→T** | **63.2%** | **42.6%** |
| **Str$^R$ Rif$^R$ 2** | 974859 | mukF | Δ3 bp | | 1.9% |
| | 1466438 | ydbA[2] | T→G | | 5.1% |
| | 2259416 | psuK ← / ← fruA[3] | +AA | | 3.4% |
| | 3150816 | yghA → / ← exbD | C→T | | 6.2% |
| | 4348862 | dcuB ← / ← dcuR[4] | IS5 +4 bp | 12.2% | 14.2% |
| | 4348971 | dcuB ← / ← dcuR[4] | Δ98 bp | 1.3% | 4.2% |
| | 4348988 | dcuB ← / ← dcuR[4] | IS2 +5 bp | 11.6% | 7.5% |
| | 4349082 | dcuB ← / ← dcuR[4] | IS2 +5 bp | | 1.4% |
| | 4349103 | dcuB ← / ← dcuR[4] | IS2 +5 bp | | 20.5% |
| | 4603111 | yjjP ← / → yjjQ | IS5 +4 bp | | 2.9% |
| | 4603111 | yjjP ← / → yjjQ | Δ2 bp | | 1.4% |

### SUPPLEMENTAL FIGURES

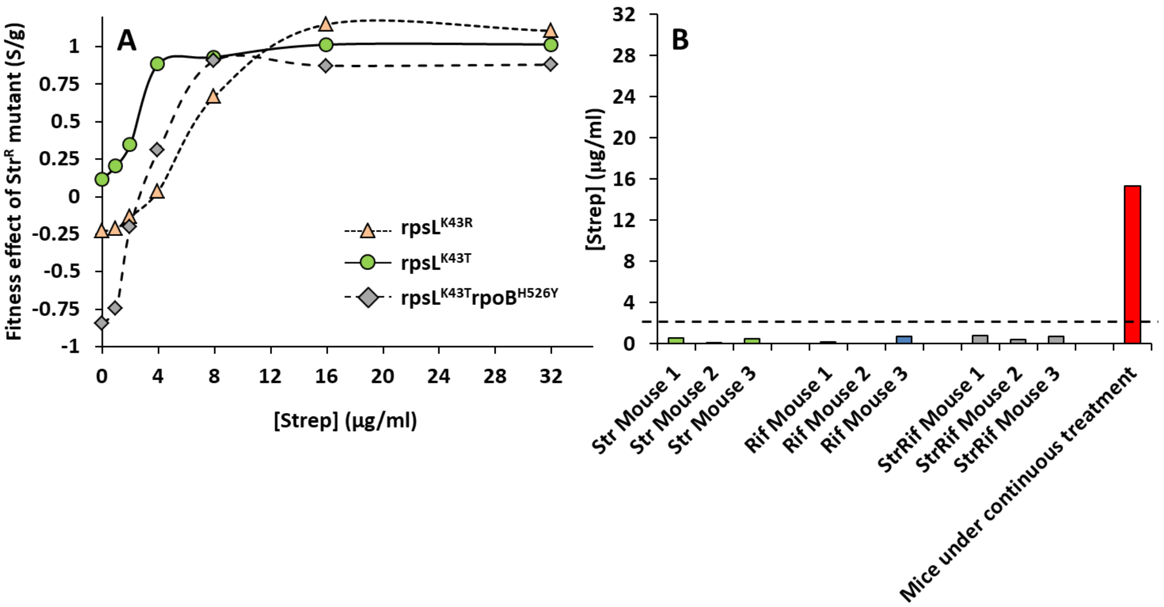


**Supplementary Figure 1 – An indirect method for streptomycin detection in fecal samples. (A)** Effect of different concentrations of streptomycin in pairwise competitions between *E. coli* MG1655 resistant to streptomycin (RpsL$^{K43T}$ or RpsL$^{K43R}$ single mutants or double mutant RpsL$^{K43T}$RpoB$^{H526Y}$) against a susceptible strain in fecal medium supplemented with LB. Fecal samples of SPF mice not treated with antibiotic were ressuspended (8 mL of PBS per gram of fecal content) and then filtered to remove the bacteria. To enrich the fecal medium with nutrients, 2ml of LB was added per 8 mL of filtered medium. Increasing concentrations of antibiotic were added to this mixed fecal medium and pairwise competitions of 1:1 between resistant and susceptible strains were performed for 24h at 37ºC. The method does not allow a reliable detection of streptomycin at concentrations below ≈2µg/ml of streptomycin. **(B)** Fecal samples of the SPF mice used in competitions were collected 4h after the gavage of *E. coli* (20h before the first time point in **Fig. 1B**) and used to detect streptomycin. Samples were filtered in 1ml PBS and 250 µl LB added and competitions between the resistant (rpsL$^{K43R}$) and susceptible strains were performed. No significant antibiotic pressure was detected in any sample.

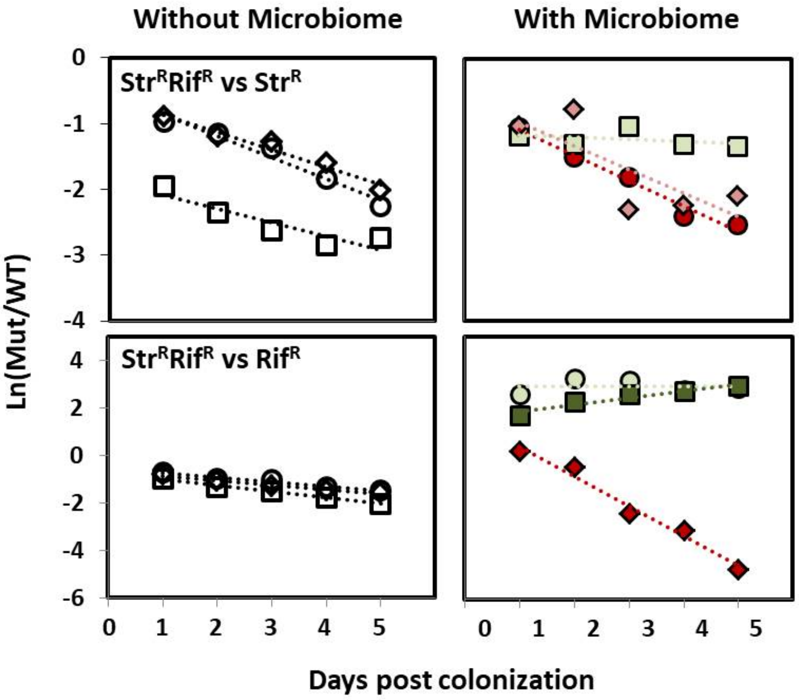


**Supplementary Figure 2 – *In vivo* fitness effects of double and single resistances are personalized in the presence of a complex microbiome.** The fitness costs of the double mutant were measured against the single Str$^R$ and against the single Rif$^R$ in germ-free (left panel) and specific-pathogen free (right panel) mice. High variance of AR fitness effects across mice is observed when competing the double against the single resistances in the presence of a microbiota. This variance is observed even when both strains are resistant to streptomycin (Str$^R$Rif$^R$ vs Str$^R$). To inquire if fitness effects were transitive, the fitness effects of these competitions were compared to the values estimated from the competitions of the single mutants against the susceptible strain (**Supplementary Table 3**).

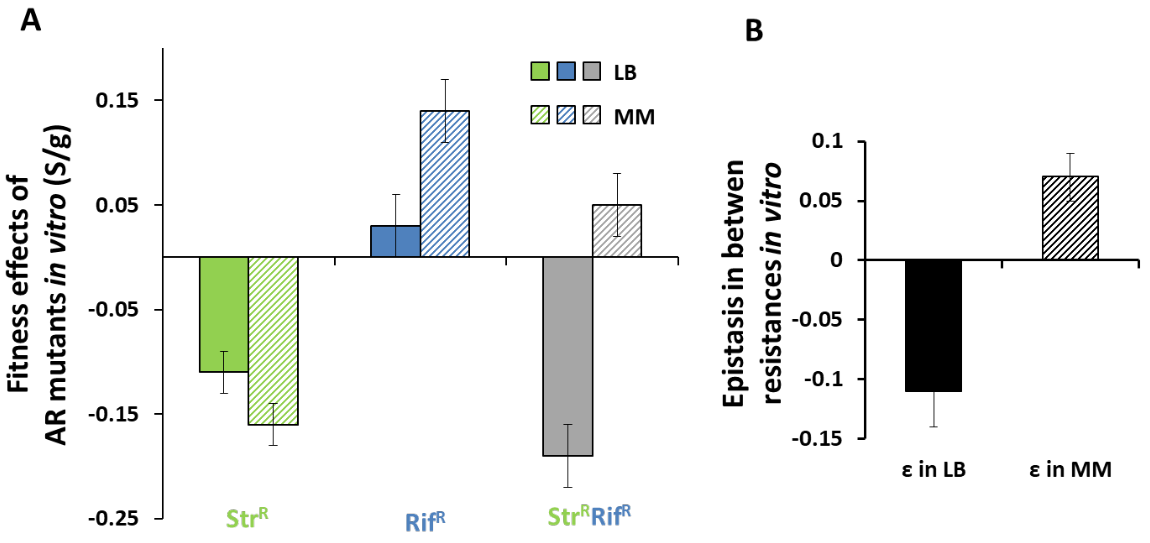


**Supplementary Fig. 3 – Fitness effects and level of epistasis *in vitro***

Fitness effects of resistances - Str$^R$ (*rpsL*$^{K43T}$), Rif$^R$ (*rpoB*$^{H526Y}$), Str$^R$Rif$^R$ (*rpsL*$^{K43T}$*rpoB*$^{H526Y}$) - in the Δ*gatZ* strains against a susceptible strain (also Δ*gatZ* background) in Luria Broth (LB) and in minimal media supplemented with 0.4% glucose (MM). Error bars represent 2*SE. Epistasis was measured assuming an additive model, and its error calculated by error propagation.

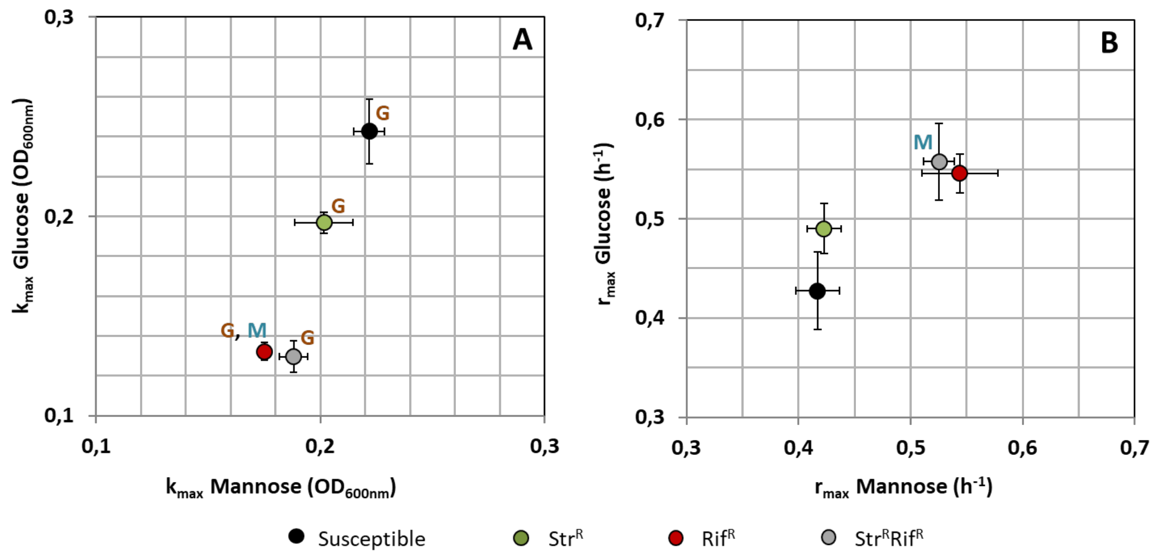


**Supplementary Figure 4 - Resistance mutations alter metabolism**. Growth was measured in minimal media supplemented either with 0.4% Glucose or 0.4% Mannose. **(A)** Carrying capacity measured as final $OD_{600nm}$ in both sugars. **(B)** Maximum growth rates in both sugars. **G** and **M** stand for significant changes in glucose and mannose, respectively (Mann-Whitney test).

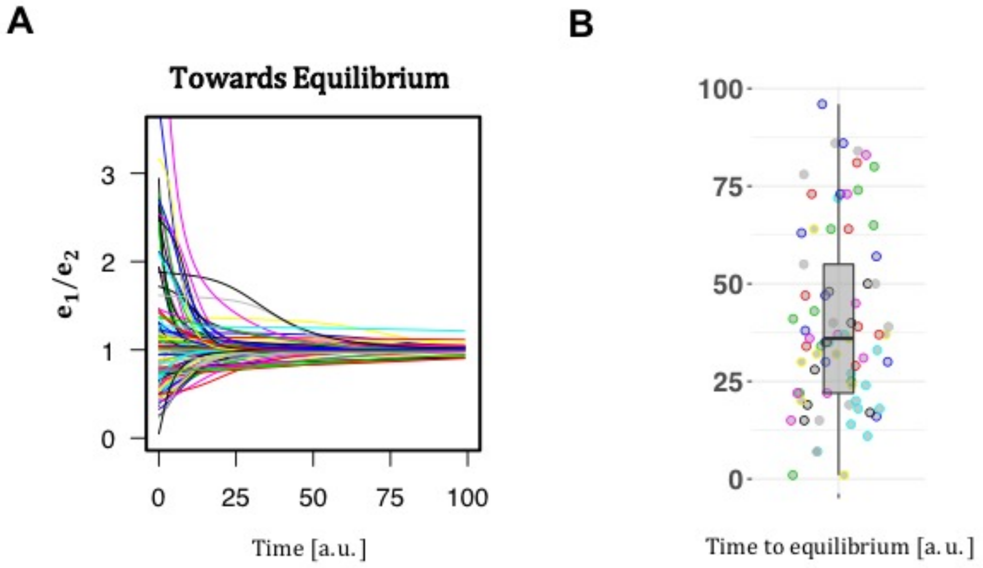


**Supplementary Figure 5 – Dynamics and time to reach equilibrium are microbiota-dependent.** Simulations of 100 different microbiotas after the perturbation over time. **(A)** The enzyme dynamics ($e_1$, $e_2$) in each of the ecosystems (different colours) are shown and are a measure of ecosystem's equilibrium. **(B)** Quantification of the time needed in panel **A** to approach equilibrium. Microbiota whose $e_1/e_2 = 1 \pm 0.01$ are considered to be at equilibrium.

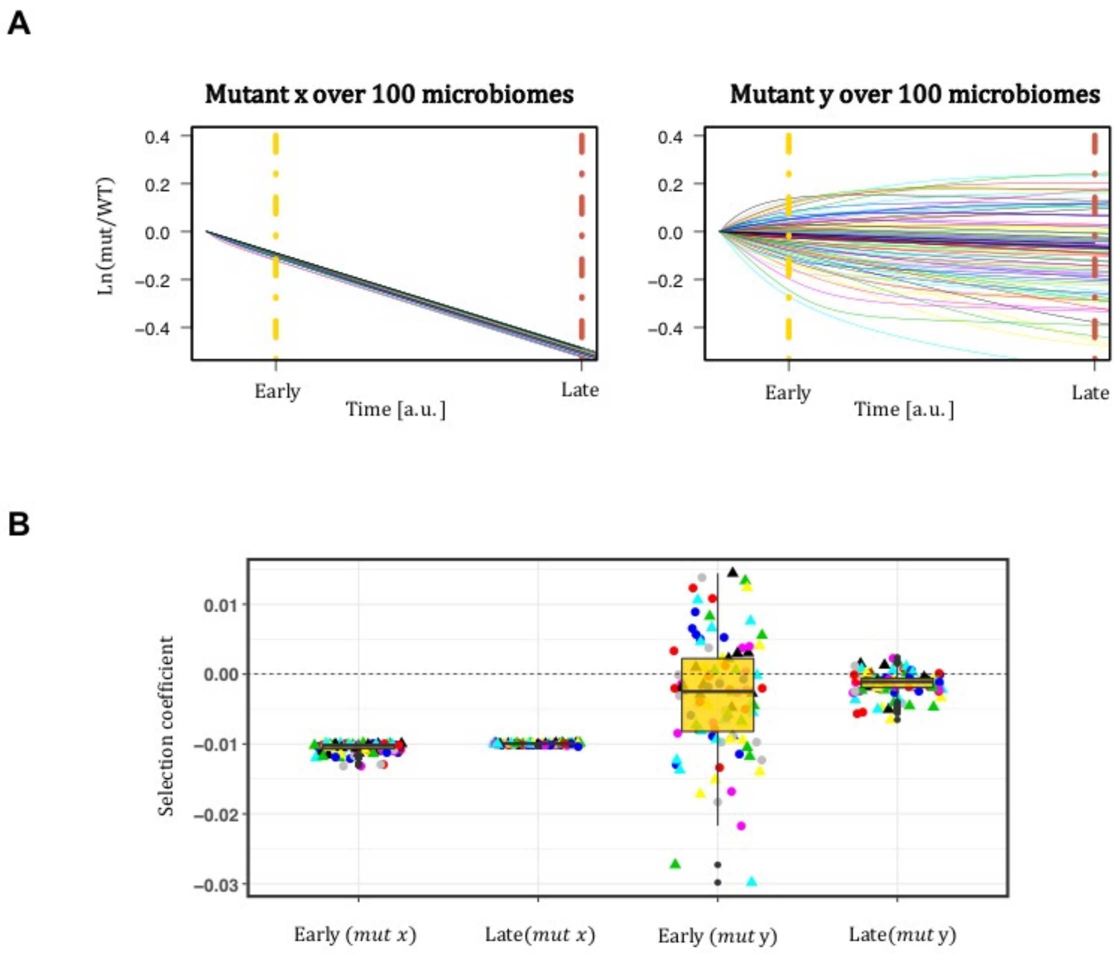


**Supplementary Figure 6 – Selection is time-dependent and depends on the level of pleiotropy of the mutation. (A)** Long-term dynamics of mutant *x* with low pleiotropy (left panel) and *y* with high pleiotropy (right panel) in each ecosystem (100 simulated microbiomes). **(B)** Quantification of the selection coefficient (slope) from panel **A**. Mutant *x*, (defined in **Figure 2)**, that represents a mutation conserving the trait ratio, shows similar fitness effect across the perturbed microbiomes, implying that pleiotropy is a necessary condition for the cost of resistance to be host-specific. Consistently mutant *y* shows dependence on the host microbiome. Over time the average cost of mutant *y* tends to the analytical prediction, which has a small value since for this mutation the presence of other species at equilibrium buffers its effect (**Fig. 2**).

### SUPPLEMENTARY TEXT

#### Resource competition model

We used a recent formalization of the MacArthur's consumer resource model capable of mimicking highly diverse communities, such as the mammalian gut microbiota[10]. To quantify the fitness effect of a mutation in the context of a complex microbiota we assumed that the mutation changes the value of the traits ($\alpha$), which code for the ability to consume each resource (see below). We first defined species by distinct values in the trait space. Then, we analytically derived the selection coefficient of a deleterious mutation, occurring within the wild-type population of a species, which reduces the trait values, both in the absence or presence of microbiota (intraspecies and interspecies interactions, respectively).

If we represent the set of resources provided to the ecosystem by $S_j$ where $j = 1, \dots, P$ and the set of species by $X^{(i)}$where $i = 1, \dots, M$, then each species is defined by its ability of consuming and processing the resource types: $X^{(i)} := (\overrightarrow{\alpha^{(i)}}, \overrightarrow{\gamma^{(i)}})$ where $\overrightarrow{\alpha^{(i)}} = (\alpha_1^{(i)}, \dots, \alpha_P^{(i)})$ is the vector of consumption rates of the $P$ resources and $\overrightarrow{\gamma^{(i)}} = (\gamma_1^{(i)}, \dots, \gamma_P^{(i)})$ is the corresponding yield (nutrient to mass transformation rate). For simplicity, the latter is assumed to be species and substrate independent ($\gamma_j^{(i)} = \gamma\ \forall i, j$). It has been showed that the dynamics of species abundances follows[10]:

$$\frac{dn^{(i)}}{dt} = n^{(i)} \left( \sum_{j=1}^{P} \frac{\gamma \alpha_j^{(i)} S_j}{\sum_{k=1}^{M} n^{(k)} \alpha_j^{(k)}} - \delta \right), \text{(i=1, ..., } M\text{)} \quad (1)$$

Where $n^{(i)}(t)$ is the density of species $i$, $\alpha_j^{(i)}$consumption rate of substrate $j$ by species $i$, $S_j$ constant substrate $j$ supply, $\gamma$ yield and $\delta$ microbial death rate.

The dynamical system (1), which holds for different growth functions (e.g. linear and the Monod function), assumes that metabolic reactions happen faster than cell division (i.e. it assumes a timescale separation ($\frac{dS_j}{dt} = 0$)). Therefore, the $S_j$ parameter in equation (1) is time-independent and represents the constant supply rate of resource $j$. For the sake of simplicity, we focused on only two resources ($S_1, S_2$), however the general qualitative results (i.e. microbiota-dependent fitness cost) hold under a higher number of resources. The dynamics of species abundances in an ecosystem with two resources is:

$$\frac{dn^{(i)}}{dt} = n^{(i)} \cdot \left( \frac{\gamma \alpha_1^{(i)} S_1}{\sum_{k=1}^{M} n^{(k)} \alpha_1^{(k)}} + \frac{\gamma \alpha_2^{(i)} S_2}{\sum_{k=1}^{M} n^{(k)} \alpha_2^{(k)}} - \delta \right) \tag{2}$$

In the context of our experimental design, the microbial ecosystem is initially at a stable equilibrium (i.e. before antibiotic treatment). Assuming metabolic trade-off ($\alpha_1^{(i)} + \alpha_2^{(i)} = E, \forall\, species\ i\ \in M$, with $E = 1$ without loss of generality) the conditions for stable coexistence of a set of M species composing the microbiota is

$$\exists (i,k) \in M: \alpha_1^{(i)} > \frac{S_1}{S_1 + S_2} \text{ and } \alpha_2^{(k)} > \frac{S_2}{S_1 + S_2} \tag{3}$$

The densities of the species at the equilibrium satisfy: $\sum_{i \in M} n_{eq}^{(i)} \cdot \alpha_1^{(i)} = \frac{S_1}{\delta}$ and $\sum_{i \in M} n_{eq}^{(i)} \cdot \alpha_2^{(i)} = \frac{S_2}{\delta}$. Let us define the time dependent variables $e_1$ and $e_2$:

$$e_1(t) := \sum_{i \in M} n^{(i)}(t) \cdot \alpha_1^{(i)}$$
$$e_2(t) := \sum_{i \in M} n^{(i)}(t) \cdot \alpha_2^{(i)} \tag{4}$$

whose dynamics represent the functional units that characterize the environment over time and can be thought of as enzymes dynamics.

We quantified the selection coefficient of a mutation within this environment by taking a focal species wild type (WT) with phenotype ($\alpha_1^{WT}, \alpha_2^{WT}$) and introducing a mutant which differs from the WT by a small difference in the trait values:

$$\alpha_1^{mut} = \alpha_1^{WT} + \Delta_1$$

$$\alpha_2^{mut} = \alpha_2^{WT} + \Delta_2$$

Assuming that $\Delta_1 + \Delta_2 < 0$, then implies that the mutant has impaired general fitness ($\alpha_1^{mut} + \alpha_2^{mut} < \alpha_1^{WT} + \alpha_2^{WT}$) compared to its parental strain. The value of its relative fitness ($s := \frac{d}{dt}\left(ln\left(\frac{n^{mut}}{n^{WT}}\right)\right)$) is what we calculate next.

In the absence of inter-species competition ($M = 0$):

$$s_0(\mathrm{t}) = \frac{S_1\Delta_1}{n^{WT}(\mathrm{t})\alpha_1^{WT} + n^{mut}(\mathrm{t})\alpha_1^{mut}} + \frac{S_2\Delta_2}{n^{WT}(\mathrm{t})\alpha_2^{WT} + n^{mut}(\mathrm{t})\alpha_2^{mut}}$$

For a deleterious mutation appearing in a stable WT population $\left(n^{mut}(t_0) \ll n^{WT}(t_0) = \frac{S_1+S_2}{\delta}\right)$ selection is constant,

$$s_0 \cong \delta \cdot \left(\frac{S_1}{S_1 + S_2} \cdot \frac{\Delta_1}{\alpha_1^{WT}} + \frac{S_2}{S_1 + S_2} \cdot \frac{\Delta_2}{\alpha_2^{WT}}\right) \tag{5}$$

In the case of migration of the two types in 1:1 proportion into an empty environment, after the saturation of the system $\left(n^{mut}(t_0) \cong n^{WT}(t_0) \cong \frac{K_{max}}{2} = \frac{1}{2}\frac{S_1+S_2}{\delta}\right)$ selection is

$$s_0 \cong \delta \cdot \left(\frac{S_1}{S_1 + S_2} \cdot \frac{\Delta_1}{\alpha_1^{WT} + \frac{\Delta_1}{2}} + \frac{S_2}{S_1 + S_2} \cdot \frac{\Delta_2}{\alpha_2^{WT} + \frac{\Delta_2}{2}}\right)$$

and for small mutations ($\Delta_j << \alpha_j^{WT}$) becomes equivalent to equation (5). Thus negative selection acting on small effect mutations is well approximated by the linear (time-independent) function (5).

In the presence of other species of the microbiota ($M$), selection is given by:

$$s_M(t) = \frac{S_1\Delta_1}{e_1(t) + n^{mut}\alpha_1^{mut}} + \frac{S_2\Delta_2}{e_2(t) + n^{mut}\alpha_2^{mut}}$$

(6)

When the microbiota $M$ is at equilibrium$\left(e_1^* = \frac{S_1}{\delta}, e_2^* = \frac{S_2}{\delta}\right)$:

$$s_{M^*} = \delta(\Delta_1 + \Delta_2) \tag{7}$$

This holds both if the mutant is generated within a species present in the microbiota ecosystem ($WT \in M$) or if the WT and the mutant invade a stable ecosystem in relatively small numbers ($n^{mut}(t_0), n^{WT}(t_0) \ll \sum_k^M n^{(k)}$) (e.g. our experimental design).

The fitness effect of a deleterious mutation is additive in its effect on the two traits, constant and independent of the microbiota composition, whenever at least two species coexist in equilibrium.

Next, we compare selection in the absence (5) and in the presence (7) of a microbiota.

1) If the $WT$ has optimal phenotype with respect to the environment ($\alpha_1^{WT} = \frac{S_1}{S_1+S_2}, \alpha_2^{WT} = \frac{S_2}{S_1+S_2}$), the microbiota has no effect on the mutant fitness:

$$s_0 = s_{M^*} \ \forall \text{ mutants}$$

2) If the WT does not have this property but the mutant conserves the trait ratio, the microbiota has no effect on its fitness:

$$s_0 = s_{M^*} \Leftrightarrow \frac{\alpha_1^{mut}}{\alpha_2^{mut}} = \frac{\alpha_1^{WT}}{\alpha_2^{WT}}$$

3) More generally, under a stable microbiota, the cost of a mutation can be buffered ($s_0 < s_{M^*}$) if:

$$\frac{\alpha_1^{mut}}{\alpha_2^{mut}} < \frac{\alpha_1^{WT}}{\alpha_2^{WT}} \text{ and } \alpha_1^{WT} < \frac{S_1}{S_1 + S_2}$$

or

$$\frac{\alpha_1^{mut}}{\alpha_2^{mut}} > \frac{\alpha_1^{WT}}{\alpha_2^{WT}} \text{ and } \alpha_1^{WT} > \frac{S_1}{S_1 + S_2} \tag{8}$$

as represented in Fig 2C in light red shading. Consequently when $\frac{\alpha_1^{mut}}{\alpha_2^{mut}} < \frac{\alpha_1^{WT}}{\alpha_2^{WT}}$ and $\alpha_1^{WT} > \frac{S_1}{S_1+S_2}$ the presence of the microbiota increases the cost of the mutation.

This model thus predicts that the fitness effect of a mutation is only altered by the microbiota if the mutation alters the trait ratio, which is probably a common scenario. This result is shown graphically in Fig. 2C.

In addition, the more specialized the $WT$ is, the higher the probability that the microbiome will buffer the cost of a deleterious mutation. Let us consider the case where the WT has a strong preference for resource 1, $\alpha_1^{WT} > \frac{S_1}{S_1+S_2}$ (by symmetry the same holds in the opposite scenario), then:

$$P(buffer) = \frac{\arcsin\left(\frac{\alpha_1^{WT}}{\sqrt{(\alpha_1^{WT})^2 + (\alpha_2^{WT})^2}}\right) + \frac{\pi}{4}}{\pi} \quad (9)$$

representing the light red shaded over the total shaded area in Fig. 2C. This probability is plotted in Fig 2D. In the limiting cases of full preference, the maximum probability of buffering is 75%.

To sum up, when the microbiota is at equilibrium, selection on a mutant is constant and negative, independently of the microbiota composition. The value of this cost when compared to a system where only intra-species competition occurs (e.g. germ-free mouse conditions) is however dependent on the specific position of the mutant in the trait space.

We next show that if the microbiota is perturbed, the fitness cost is no longer predicted to be constant but predicted to be microbiome composition specific. This prediction is able to recapitulate the host specific fitness effect phenomena observed in the *in vivo* competition.

**Resource competition model away from equilibrium**

Here we study the temporal effect of an out-of-equilibrium ecosystem, modeling a microbiota after a perturbation, e.g. antibiotic treatment or diet change. We show how this scenario makes both sign and strength of selection on a mutation to be microbiota dependent. The time-dependent relative fitness of a mutant in the presence of a microbiota is given by (6). Assuming the mutant to be at very low abundance compared to the total microbiota ($n^{mut}(t_0) << \sum_k^M n^{(k)}$), its selection coefficient is given by:

$$s_{\mathrm{M}}(t) = \delta(\frac{S_1 \Delta_1}{e_1(t)} + \frac{S_2 \Delta_2}{e_2(t)}) \quad (10)$$

thus dependent on the microbiota composition, as out of equilibrium, $\frac{S_1}{e_1(t)} \neq \frac{S_2}{e_2(t)}$. As a consequence, selection in a perturbed microbiota is host-specific. The condition under which a mutation will increase in frequency is:

$$s_M > 0 \Leftrightarrow \frac{S_1 \Delta_1}{e_1(t)} > -\frac{S_2 \Delta_2}{e_2(t)} \quad (11)$$

As antagonistic pleiotropy is defined as $\Delta_1 > 0$ and $\Delta_2 < 0$ and e1(t) and e2(t) are always positive, equation (11) establishes that antagonistic pleiotropy is necessary for obtaining a positive fitness effect, condition on the assumption that $\Delta_1 + \Delta_2 < 0$.

This "positive phase" is only temporal. Given sufficient time, the microbiota tends towards its equilibrium and the mutation fitness will converge to its negative value $\delta(\Delta_1 + \Delta_2) < 0$. However, this time-dependent selection is enough to generate variance across hosts when measuring the cost of a mutation in a given time window (**Fig. 3C**). In addition, the time needed for a mutant to become costly depends on the microbiota composition and its state (**Supplementary Figure 5**). In some hosts the mutation will be immediately costly, while in others the same mutation will be temporarily advantageous (increasing in frequency) to then decline in frequency at a later time (e.g. top pink line in Fig 3C).

This predicts that:

- the time in which an AR mutant can persist in the gut, in the absence of antibiotics, is variable across hosts and depends on the microbiota composition.
- evolutionary compensation can be more or less likely between individuals and compensatory mutations are expected to appear at different times as the sign of selection changes at different times between hosts.

**Numerical Simulations**

We used numerical simulations to confirm the analytical predictions and to graphically represent the results. For all the simulations we set the number of types in the microbiota to be $M = 10$ and the number of resources $P = 2$. The environment-related parameters were: $\delta = 1, S_1 = S_2 = 0.5$ such that the resulting carrying capacity of the system $\left(\frac{S_1+S_2}{\delta}\right)$ is 1, with no loss of generality. $m = 100$ microbiota ecosystems were considered, with trait values normally distributed around the resource supply and under the assumption of metabolic trade-off. In practice, we took $\alpha_j^{(i)} \sim N(\mu = 0.5, \sigma = 0.25)$ with $j = 1,2 \ \forall \ i = 1, \dots, M$ and normalized such that $\alpha_1^{(i)} + \alpha_2^{(i)} = 1$. Multispecies coexistence is ensured by satisfying the conditions in (3). The *mean shift* algorithm was used to cluster linearly the "types" into species with minimum distance between clusters set to 1%. The *WT* strain is arbitrarily assigned phenotype (0.6, 0.4). Equation (10) shows that selection within a microbiota does not depend on the WT phenotype itself but on the mutation effect Δ only. Thus, choosing different WT phenotypes would change the dynamics but would not affect our conclusions on selection. Mutants *x, y* and *z* are taken by solving (5) such that their relative fitness in the absence of other species is $\cong -1\%$ and they are chosen as representation of the three possible outcomes of competition. More precisely:

- *Mutant x* has phenotype (0.59403, 0.39602). It is impaired in both traits such that their ratio is the same as the WT (3/2).
- *Mutant y* has phenotype (0.6190675, 0.38), it grows better on $S_1$ but worse on $S_2$.
- *Mutant z* has phenotype (0.55351, 0.425), it grows better on $S_2$ but worse on $S_1$.

As function of their trait ratio and equation (8), the relative cost of mutants *x, y* and *z* is equal, buffered or amplified by the presence of a stable microbiota respectively. **Fig. 2C** shows graphically the latter results for our focal mutants x, y and z and for any other mutant with a small mutation effect.

To mimic our experimental design and to study selection in an ecosystem out of equilibrium, we perform a perturbation for a short time window (*Δt=10 [a. u.]),* introduce the focal strains and then study the relative dynamics. We assume that the antibiotic treatment can affect the environment both by directly killing the bacteria and by changing the metabolism of the host. We simulate the first type by assigning to each species an antibiotic sensitivity coefficient (*AbS*), drawn randomly from the *[0, 1]* interval. This contributes to increase the death rate in a species-specific manner:

$$\frac{dn^{(i)}}{dt} = n^{(i)} \cdot \left( \frac{\gamma\alpha_1^{(i)} S_1}{\sum_{k=1}^{M} n^{(k)}\alpha_1^{(k)}} + \frac{\gamma\alpha_2^{(i)} S_2}{\sum_{k=1}^{M} n^{(k)}\alpha_2^{(k)}} - \delta \cdot (1 + AbS^{(i)}) \right).$$

The second type of perturbation acts at the host level and affects the resource supply proportion. The "stressed" host has resource intake:

$$S_1^{new} = S_1 \pm \Delta_S \; with \; \Delta_S \sim Norm(\mu = 0, \sigma = 0.25) \text{ and } S_2^{new} = 1 - S_1^{new}.$$

After the perturbation, all the parameters are set to the initial values (before the perturbation) and WT and mutant are introduced in the system with equal proportion and in relatively small numbers ($n_0^{WT} = n_0^{mut} = \frac{1}{100} \cdot K_{max}$). Next, we follow the dynamics and compute the relative cost across the tested microbiomes. *Microbiota a* and *Microbiota b* shown in **Fig. 3A** are taken from the 100 tested ones as examples of opposite scenarios where the system is unbalanced towards consuming $S_1$ and $S_2$ respectively. For representation purposes, the shapes' size is proportional to the abundances of each species ($log_{10}$).
